# Host type governs influenza evolutionary strategy across reservoir and spillover hosts

**DOI:** 10.64898/2026.09.15.751820

**Authors:** Maria A. Maltepes, Alexey Markin, Stephen Shank, Jordan T. Ort, Jared Sabre, Lambodhar Damodaran, Grant Park, Kathryn Kistler, Tavis K. Anderson, Louise H. Moncla

## Abstract

Despite its high propensity for host switching, the evolutionary mechanisms underlying influenza host adaptation remain unclear. H3Nx influenza viruses are uniquely generalist, with long-term lineages that circulate in avian, human, swine, equine, and canine hosts. Using 13,295 H3Nx sequences, we quantified host-specific adaptive evolution, and developed a pipeline to map reassortment events onto trees with measures of statistical uncertainty. We find that while H3Nx viruses in mammals undergo adaptive evolution in HA and NA, viruses in birds experience very little directional selection. Instead, avian lineages exhibit high rates of reassortment, frequently generating novel reassortant lineages that persist transiently and turn over rapidly. 29.8-47.4% of all avian reassortant lineages are purged within the first year of circulation, and reassortment shows no fitness benefit in birds. In contrast, reassorted lineages in swine are more likely to persist long-term, suggesting that reassortment in swine may be broadly beneficial. Segment-specific reassortment patterns were also distinct between avian and mammalian viruses, with NA reassorting more frequently than expected in birds, but less frequently than expected in swine. Reassortment events are enriched between mammalian, but not avian, host switches, suggesting that reassortment may be most beneficial for mediating host switches among mammalian species. Together, our data suggest that host differences drive fundamentally different evolutionary outcomes for influenza viruses, transitioning from reassortment-dominant evolution in their avian reservoir, to varying degrees of adaptation upon establishment in mammals.

## Introduction

Influenza A viruses (IAVs) have a high propensity to host-switch, posing persistent challenges for human, animal, and wildlife health (Taubenberger and Kash 2010). However, there are very few natural examples of viruses that have crossed species barriers and been well-sampled in both the reservoir and novel hosts. H3Nx influenza viruses are an unusually generalist subtype that circulate enzootically in global wild bird populations (Yang et al. 2025; Yoon et al. 2014) and have established decades-long circulating lineages in humans, swine, canines, and equines. From these established lineages, additional spillovers into humans, swine, felines, camels, mink, seals, and donkeys have occurred, providing a case study on how circulation in distinct hosts impacts the fundamental forces that shape influenza virus evolution (Trovão et al. 2024; Crawford et al. 2005; Song et al. 2008; Wasik et al. 2025; Venkatesh et al. 2020; Yondon et al. 2014; Kuchinski et al. 2025; Yang et al. 2018; Song et al. 2011; Le Sage et al. 2026; Webby et al. 2000).

Adaptation to a distinct host environment can be aided by two major evolutionary forces acting on IAV’s eight-segmented genome: mutation and reassortment. Reassortment is the only mechanism analogous to recombination in influenza viruses, permitting the instantaneous acquisition of whole gene segments during co-infection. While accumulation and selection of adaptive mutations is incremental, reassortment can rapidly introduce advantageous or purge deleterious genotypes within a population (Steel and Lowen 2014). Reassortment has been linked to the emergence of at least 3 naturally occurring human pandemics, and is thought to facilitate host switching by bringing together novel gene segments to overcome host barriers (Ma et al. 2016; Furuse et al. 2010; Nelson et al. 2008; Lindstrom et al. 2004). Reassortment within avian hosts is thought to occur frequently and freely among co-infecting, homologous strains (Dugan et al. 2008; Marshall et al. 2013). In contrast, reassorted progeny from divergent parental strains can be constrained by segment mismatch in non-avian hosts (Ganti et al. 2021; White and Lowen 2018). Reassortment is likely impacted by host-specific differences in ecology, prevalence, and virologic constraint (Lowen 2017), but is understudied due to methodologic limitations.

Reassortment inference frequently relies on comparing segment phylogenies to identify incongruence. Common approaches include constructing tanglegrams, which require subjective interpretations (de Vienne 2019; Ovadia et al. 2011), and classifying sequences into genotypes as proxies for reassortment (Youk et al. 2023). Alternatively, Bayesian models enable inference of the underlying reassortment network, but are computationally intensive and are generally limited to small datasets (Müller et al. 2020). To address these limitations, TreeSort was developed and validated to accurately detect reassortment on large influenza datasets (Markin et al. 2025). TreeSort uses the molecular clock signal in the evolution of individual gene segment trees to identify recent and ancestral reassortment events. The algorithm then maps reassortment events onto a reference tree, and provides point estimates of reassortment rates (i.e., the expected number of reassortment events per year) and associated mappings, enabling large-scale reassortment mapping along with information on host species.

Here, we leveraged the broad host range of H3Nx viruses to examine how reassortment and selection jointly impact viral evolution and host specific adaptation. We developed a pipeline to parallelize TreeSort measurements to quantify reassortment along with measures of uncertainty, and measured reassortment and directional selection in each host. We uncover patterns of adaptive evolution and reassortment that vary substantially among species, and show that reassortment reflects host-dependent patterns of segment mixing. Our data suggest differential fitness effects of reassortment across hosts, with rapid lineage turnover in birds, but longer-term persistence and fitness benefits in swine. Finally, we show that reassortment is enriched on branches leading to host switches between mammals, but not between birds, suggesting that reassortment is particularly beneficial for mediating mammal to mammal cross-species transmission. These data establish reassortment as a ubiquitous generator of viral diversity that differs systematically between avian and mammalian hosts, and suggest that host differences drive fundamentally distinct outcomes for influenza virus evolution.

## Results

### H3Nx viruses have jumped hosts multiple times

H3Nx viruses have been studied extensively in humans and swine, with less work focused on how these viruses evolve across their full host range (Wasik et al. 2025; Parrish et al. 2015; Webby et al. 2000). To reconstruct the complete evolutionary history of these viruses, we curated a dataset of 13,295 non-human H3Nx sequences sampled from horses, dogs, swine, birds, and other spillover hosts between 1963 and 2024. To mitigate sampling differences between species, we subsampled sequences by year, host, country, and subtype to generate a dataset of 5,023 full genomes among all non-human hosts. Human seasonal H3N2 viruses were included separately, subsampled by year and country, generating a dataset of 1,078 full human influenza genomes sampled from 1968-2023. Three human spillover H3N8 viruses of avian-origin were also included.

Merging these subsampled datasets brought the final dataset size to 6,104 full H3Nx genomes spanning 9 NA (N1-N9) subtypes. We inferred time-resolved phylogenies for each of the eight genomic segments, revealing host jumps that have established continuously circulating H3Nx viruses in North American avian, Eurasian avian, equine, canine, human, and swine populations (Figure 1A, Supplementary Figure 1). These host switch events match known patterns of spillover that have led to repeated spillbacks between humans and swine, establishment of H3N8 equine and canine lineages, and the recent establishment of canine H3N2 viruses from birds (Crawford et al. 2005; Song et al. 2008; Webby et al. 2000; Sharma et al. 2022; Zeller et al. 2024; Rajão et al. 2015) (Supplementary Figure 2). We recapitulate a known spillover of H3N8 viruses from birds to horses, which established the H3N8 equine influenza lineage that still circulates as of 2026. Spillover of these viruses to canines established a now-extinct canine H3N8 lineage that circulated from 1999 to 2016 (Wasik et al. 2023). In 2004, H3N2 influenza viruses spilled directly from birds to dogs, establishing the extant canine H3N2 lineage (Wasik et al. 2025). Human H3N2 viruses are descendants of the 1968 H3N2 pandemic strain. Notably, the continuously circulating swine lineages across the HA tree descend from human introductions, representing multiple independent spillover events that established distinct, endemic swine lineages now globally distributed across multiple clades (Nelson and Vincent 2015; Nelson et al. 2014). For the purpose of comparing evolution across species, we delineated host and geographic lineages (see Methods for details), and built independent trees for each, providing the framework for subsequent analyses of reassortment, adaptive substitution rates, and host-switching.

**Figure 1:**
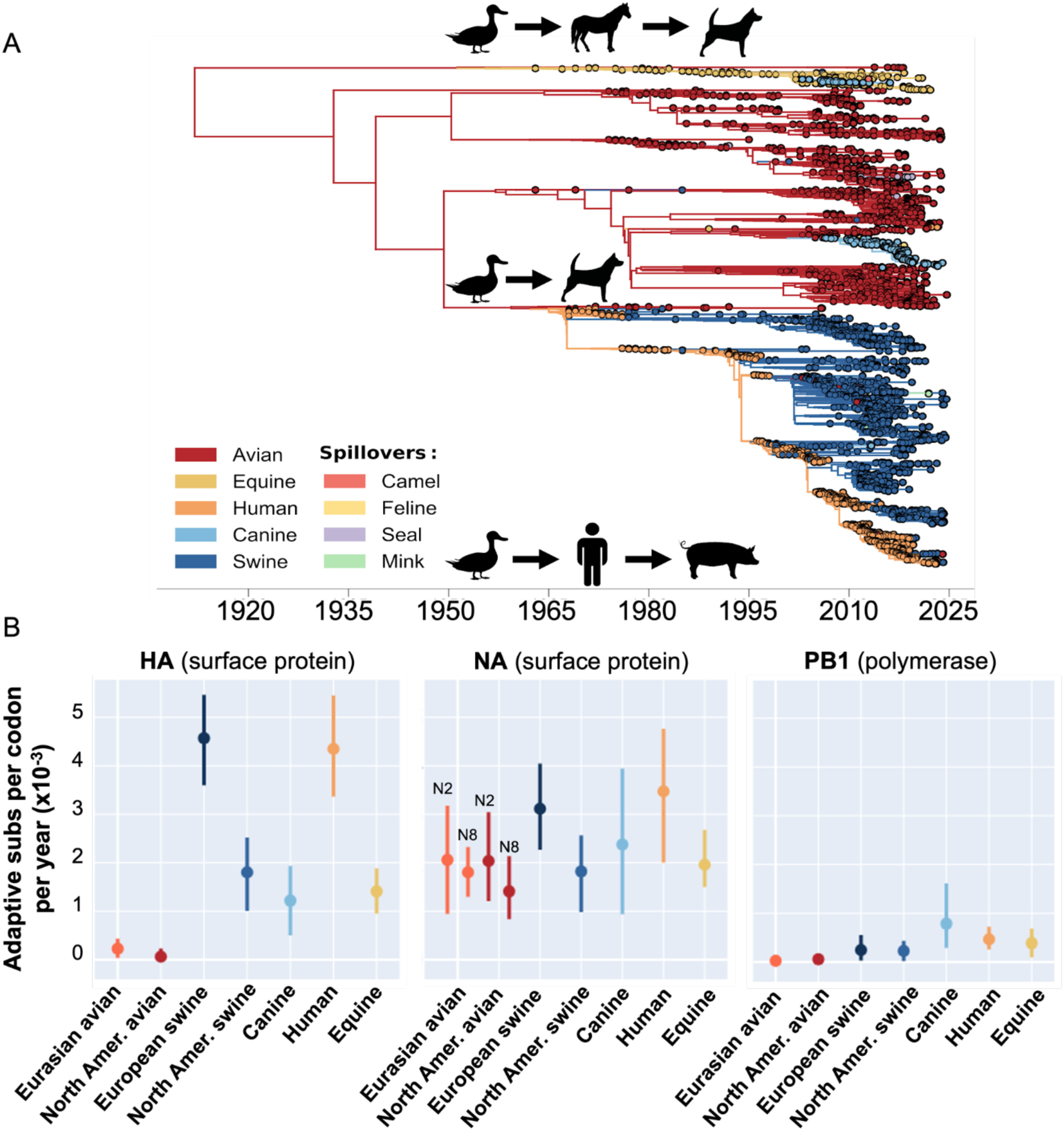
H3Nx viruses have jumped hosts multiple times, where host clades exhibit varying levels of selection. ***A)*** The HA phylogeny (N=6,104) is colored by host type, where each leaf is a unique strain and each internal node is an inferred ancestor. Notable host switches are highlighted. The 95% confidence interval for the TMRCA estimates are between 1910-05-22 – 1917-09-14. ***B)*** Rates of adaptation (adaptive substitutions per codon per year * 10^−3^) for each host are shown for the genes encoding the surface proteins HA and NA, as well as the polymerase gene PB1. Significant differences between host and genes are represented by estimates that fall outside of another estimate’s error bars.

### Adaptive substitution rates vary by host

Human seasonal influenza viruses evolve through yearly selective sweeps, in which strong, directional selection fixes HA genes that mediate immune escape (Rambaut et al. 2008). Differences in the strength of selection between species could arise from differences in life span, vaccination patterns (e.g., in horses and swine), and the frequency of repeat infections (Ferguson et al. 2003; Wille and Holmes 2020; Petrova and Russell 2018). To determine the strength of selection across each host lineage, we performed a modified McDonald-Krietman (MK) test on two surface glycoproteins (the hemagglutinin (HA), neuraminidase (NA)), and a highly conserved polymerase gene (the polymerase basic 1 (PB1)). While HA and NA are exposed to host-specific receptor and immune factors and may undergo selection, PB1 is the most conserved, and less likely to experience strong, host-specific selection. This test calculates an adaptive substitution rate by measuring the rate of nonneutral substitution accumulation over time that exceeds a neutral expectation, and was developed specifically for temporally-sampled viral genomes (Kistler and Bedford 2023; Bhatt et al. 2011). Because this test assumes that viruses are co-circulating and competing with each other to at least some degree, we subsetted viruses by host and geography for analysis (see Methods).

Adaptive substitution rates varied substantially between gene segments, with the highest adaptive rates in the HA and NA surface glycoprotein genes. Adaptive substitution rates were largely similar across all host groups for PB1, in line with previous findings suggesting limited direction selection on this polymerase protein (Kistler and Bedford 2023). Human seasonal H3N2s and European swine viruses exhibited the highest rates of positive selection in HA, followed by canine, equine, and North American swine lineages. Differences in adaptive rates between North American and European swine groups could arise due to regional variation in farming practices, swine movement, and vaccination coverage, or from the distinct evolutionary histories of these lineages. We found no detectable signal of adaptive evolution in HA among Eurasian and North American avian lineages, with adaptive rates equivalent to those measured in PB1. Adaptive rate estimates for NA were moderately elevated, but overlapped across all host lineages, with no host lineage exhibiting significantly higher rates relative to all other hosts (Figure 1B, Supplementary Figure 3). These data indicate that surface protein genes experience distinct selection patterns across species, with very little directional positive selection on HA in birds, and moderate selection in swine, canine, and equine lineages.

### Reassortment rates vary by host and geography

In wild aquatic birds, low pathogenicity avian influenza viruses are thought to evolve frequently via reassortment, a process in which genomic segments from co-infecting virions mix within an infected host cell (Dugan et al. 2008; Wille et al. 2013). Reassortment in wild birds is hypothesized to facilitate immune escape by mediating the continual generation of novel subtypes, which may reduce the necessity of antigenic selection (Roche et al. 2014). Influenza viruses also reassort frequently in swine, with reassortment known to be associated with changes in phenotype that may mediate interspecies transmission (Smith et al. 2009; Lycett et al. 2012; Khiabanian et al. 2009; Thomas et al. 2024). We next aimed to determine whether reassortment rates and patterns varied across hosts.

Reassortment inference is frequently limited by a lack of methods that can perform robust inference across large datasets. TreeSort is an approach validated for identifying reassortment among influenza viruses (Markin et al. 2025) that designates one segment as the reference, and then systematically compares the evolutionary histories of the remaining segments to identify signatures of incongruence. To account for statistical uncertainty, we developed a pipeline in which TreeSort is run 1000 independent times, and a summary reassortment tree is generated in which each reassortment event (and the segments involved) is annotated with a support value. We then produce summary trees retaining only high-support reassortment events that occur in at least 95% of replicates (Figure 2B, Figure 2C, Supplementary Figure 4).

**Figure 2:**
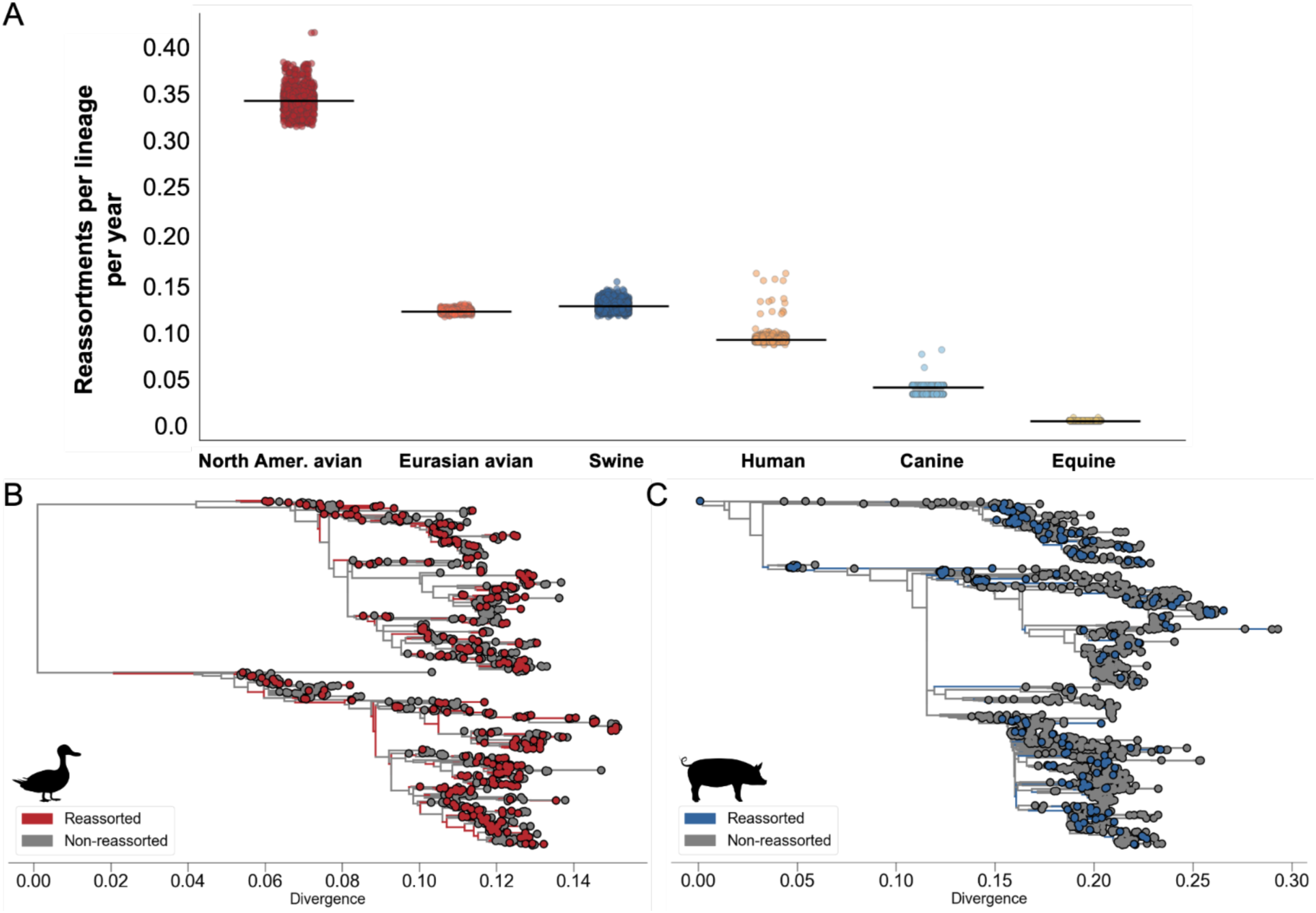
Reassortment rates vary across hosts and regions. ***A)*** Reassortment rates (reassortments per lineage per year) were calculated for each H3Nx host group. Each point represents a reassortment rate calculated for an individual TreeSort replicate. All 1000 replicates are shown, with the mean value indicated with a line. Though more high-support reassortments were called in Eurasian avian viruses (415 reassortments) than in swine viruses (307 reassortments), the lower average rate in Eurasian avian viruses reflects differences in the host-specific molecular clock rates used to calculate the rate (see Methods). Summary TreeSort trees are shown for the ***B)*** North American avian and ***C)*** swine host groups. Branches are colored to indicate reassortment events supported at >= 95%.

Reassortment rates were generally robust across TreeSort replicates. However, the mapping of reassortment events onto specific branches varied across replicates, particularly for high-reassortment hosts (avian and swine) (Supplementary Figure 8). Overall, 54.4% of inferred reassortment events met the 95% support threshold, supporting the utility of our replicate-based approach for clarifying high-support events. Using these high-support events, we calculated reassortment rates for each host clade. Host-specific reassortment rates were highest in North American avian viruses (mean ± sd: 0.3472 ± 0.0131 reassortments per year per strain), and markedly lower in Eurasian avian viruses (0.1248 ± 0.0018). While H3s are among the most prevalent subtype in North American birds, H6 and H4 subtypes dominate in European wild migratory birds, which may account for these differences (Diskin et al. 2020; Munster et al. 2007). Within North America, reassortment was most common in the Mississippi flyway, suggesting that geographic variation may also impact rates (Supplementary Figure 5), though expanded datasets are necessary to confirm these results. Mammalian hosts exhibited lower reassortment rates overall, with the highest mammal rates observed in swine (0.1303 ± 0.0052), and lower rates in human (0.0952 ± 0.0063), canine H3N2 (0.0446 ± 0.0044), and equine (0.0096 ± 0.0001) (Figure 2A) lineages. Overall, patterns of reassortment and adaptive substitutions were roughly inverse across species, with the highest reassortment rates and lowest adaptive substitution rates in birds, and low reassortment and higher adaptive substitutions in equines, canines, and humans. Uniquely, swine influenza viruses exhibited both high reassortment rates and detectable adaptive evolution in HA and NA.

To assess the sensitivity of reassortment inference to dataset size, we generated serially subsampled datasets of North American avian viruses, ranging from N=400 to N=1100 (increasing by 100 sequence increments, 5 trials each). Reassortment rates increased with dataset size, suggesting that absolute rates may be sensitive to sampling (Supplementary Figure 6). However, the relative ranking of reassortment rates was consistent across datasets, with North American avian viruses exhibiting the highest rates even when heavily downsampled (N=400). Similarly, North American avian viruses exhibited higher rates than swine despite fewer available sequences. We also found high reassortment support for a range of root-to-tip divergence values, indicating that TreeSort does not preferentially detect recent reassortments (Supplementary Figure 7A). Similarly, reassortment support showed minimal correlation with branch length, suggesting that reassortment inference is not biased towards long, divergent branches (Supplementary Figure 7B). Prior TreeSort validation has also demonstrated a limited association between reassortment rates and genomic diversity, suggesting that observed differences in reassortment frequencies are not solely driven by differences in circulating diversity across these populations (Markin et al. 2025). Together, these data suggest that the differences in reassortment rates among hosts cannot be explained purely by differences in sampling or diversity.

### Segments reassort nonrandomly

Influenza reassortment is known to be restricted by segment compatibility (White and Lowen 2018; Lowen 2017). HA and NA must be functionally compatible, while compatibility among segment packaging signals can further augment reassortment viability (Mitnaul et al. 2000; White et al. 2017; Baker et al. 2014; Essere et al. 2013; Liu et al. 2022). To determine whether segment-specific reassortment patterns differed among hosts, we quantified segment specific reassortment frequencies. We then compared these results to expectations under a null distribution generated by simulating the same number of reassortment events as observed in our actual data, but randomly sampling segments for each event with equal probability. This null thus represents the expected frequency of reassortment of each segment under a model in which reassortment is random. Given the relatively low rates of observed reassortment in human, canine, and equine lineages, we focused subsequent reassortment analyses on the avian and swine host groups for which we had sufficient statistical power. For all subsequent analyses, we report reassortment events relative to HA’s evolution.

In both avian and swine lineages, many segments reassorted nonrandomly (p < 0.05 after Bonferroni correction). Among North American avian viruses, NA and PA were significantly overrepresented in reassortment events (p = 0.007 for both), while NS and MP were underrepresented (p = 0.02 and p = 0.007) (Figure 3A). In Eurasian avian viruses, NA and NS reassortments were overrepresented (p = 0.007 and p = 0.02) and PB1, PA, and MP were underrepresented (p = 0.02, p = 0.007, and p = 0.007, respectively; Supplementary Figure 9A). Despite past evidence pointing to permissive gene segment combinations in birds (Ganti et al. 2021; Dugan et al. 2008), these patterns indicate that gene mixing is not entirely free or random among H3Nx viruses. Instead, we find evidence in both Eurasian and North American avian populations for increased NA reassortment events. Non-random reassortment could reflect functional constraints on segment compatibility or non-random co-circulation of subtypes in the avian reservoir, leading to unequal co-infection frequencies. In swine, reassortment patterns were distinct, with reassortments involving NA and PB1 underrepresented (p = 0.007 and p = 0.03) (Figure 3B). The high overall rates of reassortment in swine confirm that opportunities for co-infection and reassortment occur readily, while the strong observed linkage between HA, NA, and PB1 (which rarely reassort away from each other), could reflect that these gene pairs are selectively beneficial, or could arise from constraints in gene segment packaging that conserves these pairings (Lowen 2017; Nelson et al. 2014; Mitnaul et al. 2000; Ma et al. 2012). Because swine H3Nx viruses all descend from spillovers from humans, it is also possible that these findings reflect maintenance of these gene pairings in human-adapted H3N2 strains following spillover (Yen et al. 2011; Xu et al. 2012). While our data cannot distinguish the reasons for these patterns, they do show that reassortment patterns in both swine and avian species are non-random, and differ substantially from each other.

**Figure 3:**
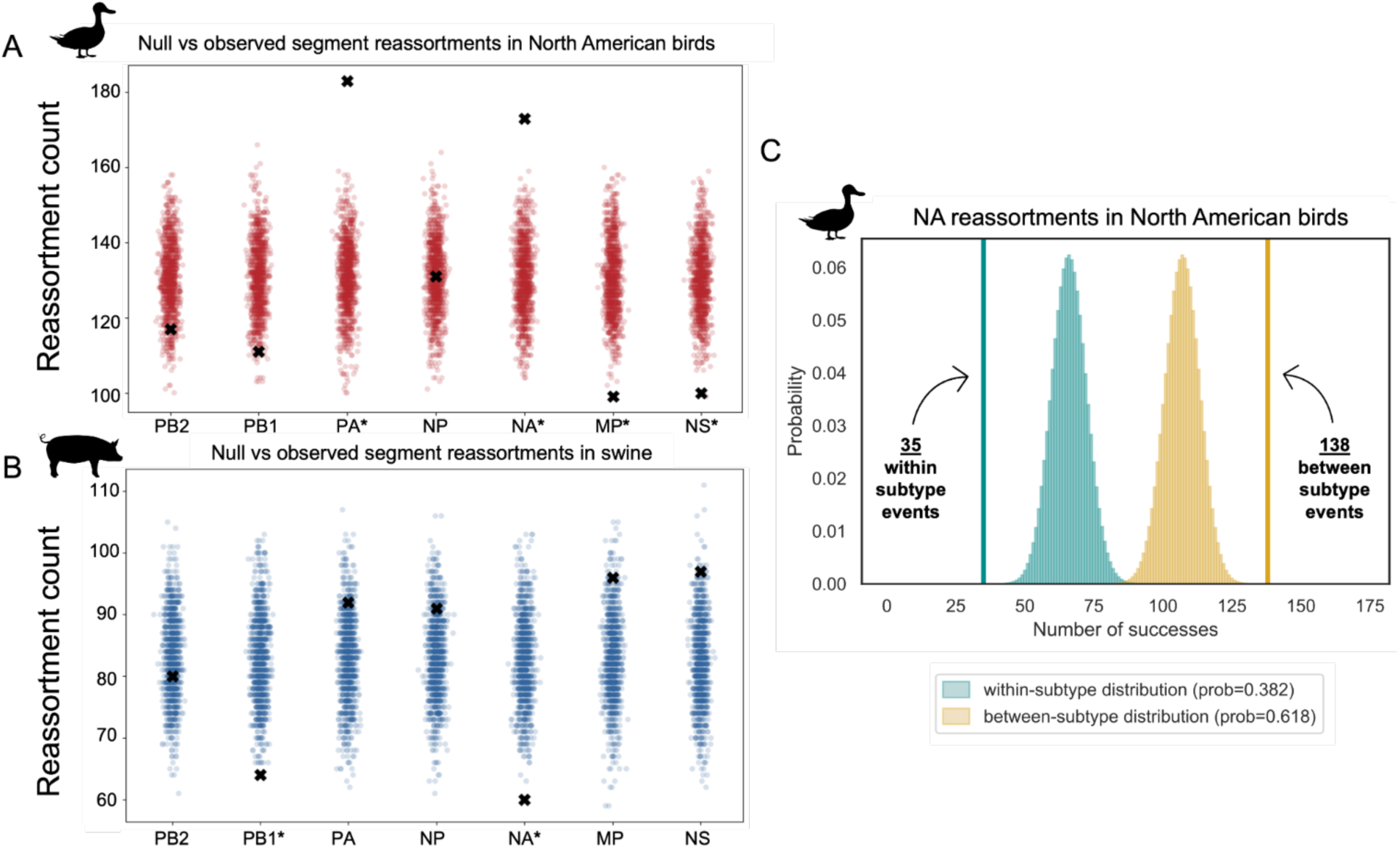
Segments exhibit host-specific compatibility with HA. Segments reassort nonrandomly relative to a null model in ***A)*** North American avian and ***B)*** swine viruses. Data points represent simulated reassortment counts for each segment under random sampling while observed counts are denoted with a black X. Asterisks indicate p-value < 0.05 after Bonferroni correction. ***C)*** Binomial testing revealed a significant excess of between-subtype NA reassortments (p-value = 3×10^−7^). Success probabilities for the binomial model were defined as the within-subtype (0.382) and between-subtype (0.618) NA reassortment probabilities. Observed within-subtype (blue, 35 events) and between-subtype (yellow, 138 events) NA reassortments are indicated by vertical lines.

### Novel NA subtypes are introduced at a rate exceeding expectations based on viral cocirculation alone

In both European and North American birds, we observed an excess of reassortments involving NA, but did not distinguish between within and between subtype events. Within-subtype reassortment events could be favored if only some combinations of HA and NA are functionally compatible, allowing within-subtype reassortments to better maintain HA:NA balance. Alternatively, if high subtype diversity is preferentially maintained in wild birds (e.g., through selective or ecological processes), then reassortment of new NA subtypes might occur more frequently. Using the frequencies of NA subtypes within our dataset, we calculated the expected frequency of within vs. between subtype reassortment events as the probability of sampling two sequences of the same, or distinct, NA subtypes from the population (Supplementary Figure 10). We then classified all observed NA reassortment events as within- or between-subtype events using a divergence-based threshold (see Methods for details). Analysis of pairwise divergence values showed clear delineation between pairwise divergence rates for within vs. between-subtype NA sequences, supporting this approach (Supplementary Figure 11, Supplementary Figure 12). We identified 35 within-subtype and 138 between-subtype reassortment events in the North American avian lineage, and 39 within-subtype and 109 between-subtype events in the Eurasian avian lineage. For avian viruses, the expected frequency of within- and between-subtype reassortments were 0.382 and 0.618, respectively for North American viruses, and 0.412 and 0.588 for Eurasian viruses. Comparing observed counts to these expected probabilities under a binomial model revealed a significant excess of between-subtype reassortments in both avian populations (Figure 3C, Supplementary Figure 9B).

Reassortment detection depends on identifying lineages that are more divergent than expected, and could be biased towards detecting between-subtype reassortment events. To estimate the robustness of our results to this bias, we conducted a sensitivity analysis to estimate the robustness of this finding to varying underdetection rates for within-subtype events. We first calculated the divergence value among all detected NA reassortment events, and then determined the fraction of within-subtype NA sequence pairs that were less divergent than the minimum detected reassortant value. Among avian viruses, 1.6-2.4% of all NA sequences had pairwise divergences that fell below the minimum divergence value of detected reassortments, suggesting the potential for TreeSort to misclassify these events as false negatives. By sequentially assuming varying rates of false negatives, we estimate that our finding of excess between-subtype reassortants is robust to an underdetection rate of 7% (for Eurasian avian) to 11% (for North American avian), substantially higher than our estimated underdetection rate (1.6-2.4%) (Supplementary Figure 13, Supplementary Figure 14). These data suggest that in both Eurasian and North American avian populations, novel NA subtypes are introduced by reassortment more frequently than expected from cocirculation alone. Furthermore, this finding is robust to a moderate degree of underdetection of within-ubtype reassortment events.

### Reassortment provides differential fitness effects across host groups

The high reassortment rate and non-random segment patterns observed in avian and swine lineages supports reassortment as a critical component of the evolutionary process in these species. To assess whether reassortment confers fitness advantages within avian and swine lineages, we used reassortant lineage persistence as a proxy for viral fitness. Selectively “fit” viruses should produce more offspring, which can be quantified by assessing how long descendant lineages persist into the future (Müller et al. 2020). Here, a reassorted lineage was defined as the longest path between a reassortment event and either a subsequent reassortment event or terminal node, with persistence measured in years of circulation. This analysis was restricted to the avian and swine host groups due to limited statistical power in the remaining mammalian hosts. While average reassortant persistence times did not differ significantly between avian and swine (Supplementary Figure 15), we observed notable differences in reassortant lineage turnover rates. North American avian reassortant lineages experience rapid turnover, with approximately 47.4% of these lineages purged within the first year of circulation. These findings align with previous, smaller-scale studies demonstrating that circulating genotypes are continuously replaced by novel genotypes created through reassortment (Macken et al. 2006). In contrast, only 29.8% and 34.7% of reassortant lineages are purged within the first year in Eurasian avian and swine viruses, respectively, indicating a much higher fraction of reassortment lineages that persist into the future.

To determine whether differences in turnover demonstrate differential fitness effects (vs. simply different reassortment rates), we compared reassortant lineage persistences to those under a null model. We shuffled reassortment events across the trees 1000 times and compared persistence times from our observed data to those calculated from these null, shuffled datasets. Persistence times of reassortant lineages showed contrasting patterns across host groups. North American avian reassortant lineages generally persisted in accordance with null expectations (Figure 4A), while Eurasian avian reassortant lineages showed decreased survival compared to the null model at later persistence times (Figure 4B). In contrast, swine reassortant lineages persisted marginally longer than expected under null expectations in the short-term (Figure 4C), indicating that reassortment in swine may confer a fitness advantage. These patterns were also recapitulated when comparing reassortant and nonreassortant lineage persistence within each host group (Supplementary Figure 16), indicating observed lineage turnover frequencies are not an artefact of tree structure.

**Figure 4:**
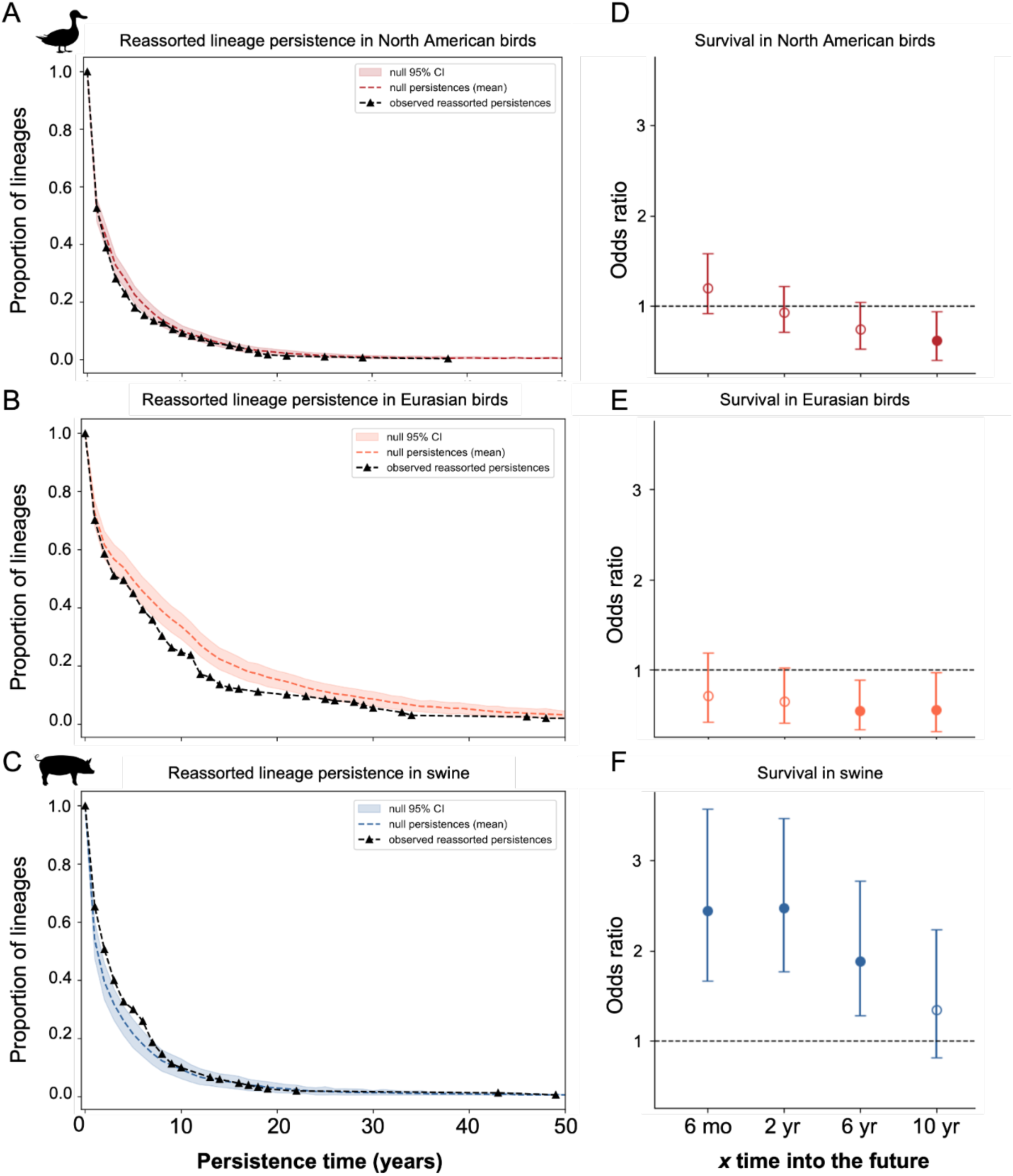
Reassortment exhibits differential fitness effects by host. Reassortant persistence analyses for ***A)*** North American avian, ***B)*** Eurasian avian, and ***C)*** swine host groups. Shaded regions indicate persistence values calculated under a null model of random reassortment. Dashed lines represent observed reassortant persistence values. Each time bin represents the proportion of reassortant lineages still in circulation at that time. ***D–F)*** Long-term survival effects of reassortment were quantified using Fisher’s exact test, where fitness was defined as a reassortment event sustaining descendants at x = 0.5, 2, 6, and 10 years into the future. Points represent odds ratios (OR) with error bars denoting 95% confidence intervals. Filled points indicate statistically significant results (p < 0.05).

We next quantified the probability that reassortant and non-reassortant lineages sustain descendants at 0.5, 2, 6, and 10 years post-reassortment using Fisher’s exact tests. If reassortment is associated with a fitness benefit, we reasoned that reassortant lineages should be more likely to sustain long-term descendants than non-reassortant lineages. In North American avian viruses, reassortment had no significant effect on long-term survival across most time points, with the exception of a marginally significant decrease in survival at *x* = 10 years (OR: 0.63; p = 0.04). The odds ratios decreased with increasing time intervals, suggesting a trend toward reduced long-term fitness (Figure 4C, left panel). This trend was recapitulated in Eurasian avian viruses, where significant decreases in survival emerged at *x* = 6 and *x* = 10 years (OR: 0.52 and 0.56; p = 0.008 and 0.046, respectively). In contrast, swine viruses showed significantly elevated survival probabilities at all time points except *x* = 10 years. Critically, at x = 0.5 and 2 years, survival effects of reassortment in swine viruses fell outside the distribution of odds ratios under a null model of reassortment (Supplementary Figure 18), suggesting that these results are unlikely caused by underlying tree topology. While we cannot exclude the potential impacts of swine population structure and turnover and immune heterogeneity on these findings, our data suggest an association between reassortment and long-term lineage persistence in swine. These results are consistent with surveillance data indicating that most novel swine genomes persist for ∼1.8 years, though dominant lineages can persist much longer (∼10 years) (Janzen et al. 2025). Together, these differences in long-term survival mirror the trends observed in the reassortant persistence analysis, pointing to a potential association between reassortment and fitness in swine, but not avian, viruses.

### Reassortment is enriched for switches between mammalian hosts

Reassortment is a key driver of pandemic virus formation, and is thought to be critical for host switching (Ma et al. 2016; Ince et al. 2013; Mehle et al. 2012; Taubenberger and Kash 2010). To determine whether reassortment was statistically associated with host switches, we classified branches in the tree as “host switch” or “not host switch” branches, and evaluated the frequency of reassortment events on each branch type. We observed a significant enrichment of reassortment on host-switching branches (OR= 5.49, p-value= 9 × 10^−7^, 95% CI=3.08, 9.81, Fisher’s exact tests) (Figure 5A), consistent with previous findings linking these processes. Among the 99 total host switches identified in the global H3Nx tree, 35 (35%) involved reassortment. Of these reassorted host switch branches, 21 occurred on internal nodes, which comprised of mammal-to-mammal (18 events: 17 human-to-swine, 1 swine-to-human), avian-to-mammal (2 events: 1 avian-to-seal, 1 avian-to-human) and mammal-to-avian (1 event: swine-to-avian) transitions (Figure 5C). Details of these reassortment events, including those occurring on terminal nodes, and the segments involved are provided in Supplementary Table 1.

**Figure 5:**
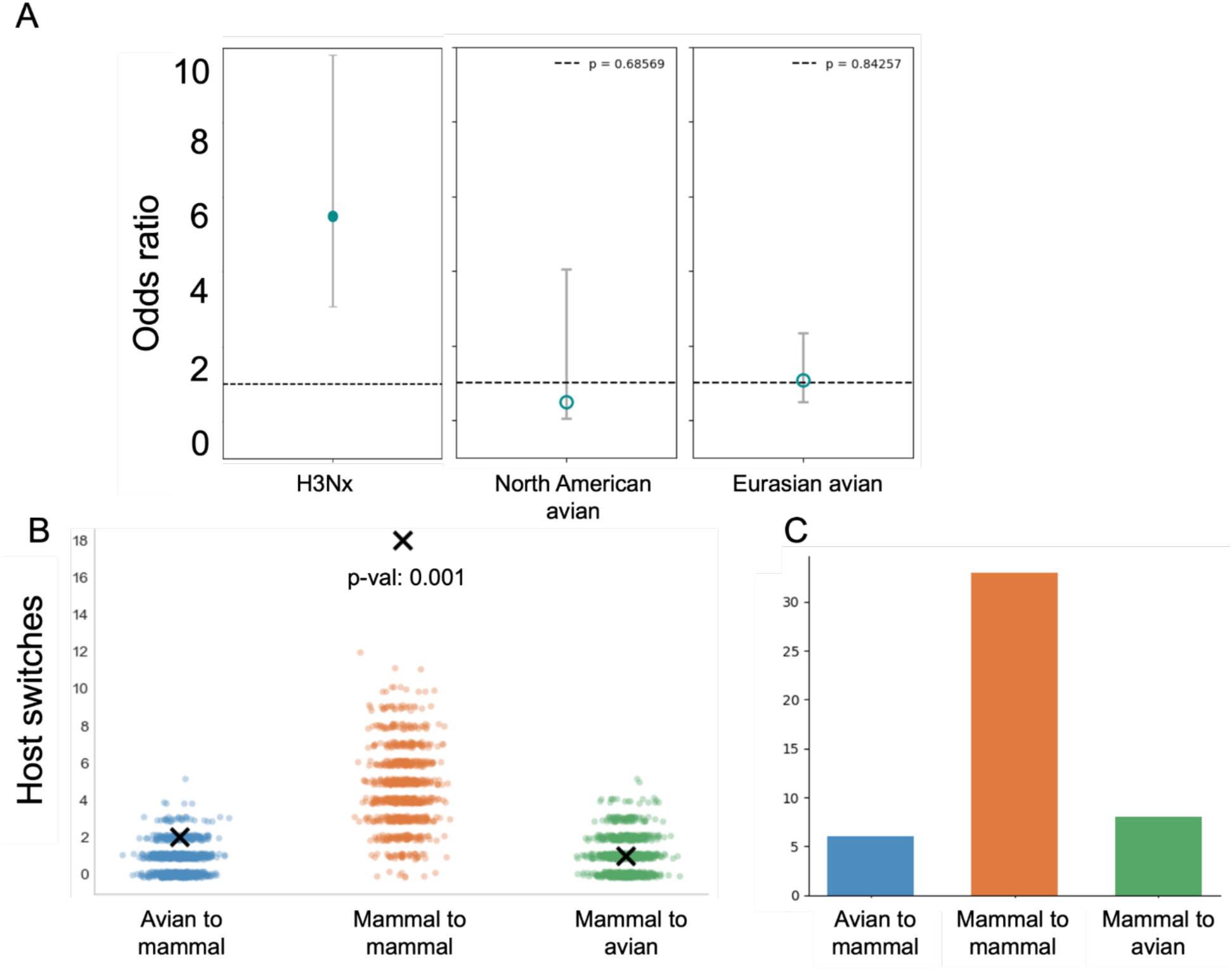
Reassortment is enriched for host-switching between mammalian hosts, but not avian hosts. Fisher’s exact test assessing whether reassorted branches are more likely to be host-switched yielded a significant association for ***A)*** all host transitions but no significant association when restricted to switches between avian orders. ***B)*** For each transition type, observed reassortment events on host-switched internal nodes (black X) are compared against expectations under a random model of reassortment (colored points). Enrichment is significant only for mammal-to-mammal transitions (p = 0.001; orange), with no significant enrichment for avian-to-mammal (blue) or mammal-to-avian (green) transitions. ***C)*** Total host-switch counts, on both internal and terminal nodes, are shown for each transition type.

To determine whether this association varied significantly across the taxonomic groups, we compared the observed frequency of reassorted host-switched internal nodes to the expected frequency under random reassortment. This analysis revealed that reassortment enrichment was taxonomically specific. Reassortment was significantly overrepresented on nodes representing mammal-mammal host switches, with the majority being human-to-swine transitions (Figure 5C). In contrast, no significant enrichment was detected for host switches between avian orders (North American avian: OR=0.49, p-value= 0.69, 95% CI= 0.058, 4.05; Eurasian avian: OR= 1.08, p-value= 0.84, 95% CI= 0.50, 2.33), which include switches between Anseriformes (ducks, geese) and Galliformes (chickens, turkeys) (Figure 5A, Figure 5B). These results suggest that reassortment may play a distinct role in mammalian adaptation compared to avian host switching. However, this analysis cannot determine the temporal relationship between reassortment and host-switching events, leaving open whether reassortment precedes or follows successful host switches.

## Discussion

We established a robust phylogenetic approach for evaluating how host-specific ecology and biology shape fundamental evolutionary processes in IAVs. Our data reveal a dichotomy in how mutation and reassortment contribute to IAV diversification across hosts. Across species, adaptive substitution rates fall along a spectrum, with the highest rates in humans, followed by other mammals, and the lowest rates in birds. Reassortment rates also vary substantially, but in the opposite direction, with the highest rates in birds, and lowest rates in canines and equines. Among the H3Nx viruses, mammalian lineages originally descend from spillovers from birds (and for swine lineages, subsequent human to swine spillovers), suggesting that influenza viruses can switch from a reassortment-dominant mode of evolution in birds to varying degrees of adaptive evolution upon establishment in new species. The variation in these evolutionary strategies across hosts may arise from a combination of ecological, biological, and epidemiologic differences between species, including lifespan, aggregation/housing, vaccination, and the degree of influenza diversity circulating in that population. These features likely interact to determine whether mutation or reassortment predominates in driving viral diversification across host species, highlighting the flexibility of these viruses to modulate the strength of adaptive evolution and reassortment. This flexibility may be one factor that enables efficient host switching and adaptation, which is uniquely common among the H3Nx viruses.

Despite decades of study, surprisingly little is still known about how influenza viruses evolve in their natural reservoir. We observe low adaptive rates in birds, which may reflect the migration and shorter lifespans of wild bird species (particularly ducks, which dominate our datasets), leading to more frequent population turnover when new immunologically naïve susceptibles enter the population (Munster et al. 2007). As the primary reservoir host, avian species also harbor extensive subtype diversity of low-pathogenicity viruses, enabling mixing with a broad diversity of subtypes that co-circulate. Under these conditions, reassortment may become the primary driver of viral evolution in avian IAVs. We find that avian reassortant lineages are short-lived and show no evidence of being selectively beneficial. The high reassortment rate coupled with rapid reassortant lineage turnover in birds is consistent with a model in which short avian lifespans and environmental durability of IAV promote a diverse reservoir that repeatedly exposes naïve hosts to novel subtype combinations (Roche et al. 2014). These data suggest that avian influenza viruses rely primarily on reassortment for their evolution, with reassortment serving as a frequent, but largely stochastic phenomenon. In contrast, the longer lifespans, biosecurity measures, and/or reduced environmental persistence of IAV in other host species may limit opportunities for reassortment. Finally, lower observed reassortment rates in other species with short life spans, like swine, may be a consequence of reduced number of subtypes that co-circulate within these species, providing fewer opportunities for reassortment to occur.

All H3Nx viruses circulating in swine reflect interspecies transmission events from humans, with subsequent evolution occurring within established, enzootic swine virus populations. However, these patterns vary substantially between lineages circulating in North America and Europe. H3N2 is geographically restricted in Europe, with widespread circulation primarily reported in Germany, Italy and the Netherlands (Brown 2013; Simon et al. 2014). The European swine lineage (H3 1970.1) is most similar to human-seasonal viruses detected in the 1970s and has circulated in European swine since (Vincent et al. 2020). In contrast, North American swine lineages are predominated by H3N2 viruses that have been seeded by recurrent human-to-swine spillover events occurring multiple times since the late 1990s (Nelson and Vincent 2015; Vincent et al. 2020), that then diversified. We find that swine influenza viruses exhibit the unique combination of high rates of reassortment and directional positive selection on HA and NA, a combination not observed in any other host group. We also find an association between reassortment and viral persistence and lineage survival in swine, pointing to a fitness advantage conferred by reassortment. Prior work has found that influenza A viruses are detected in in farmed pigs year-round, and that multiple cocirculating lineages can be detected within individual farms (Kyriakis et al. 2017; van der Vries et al. 2025), potentially facilitating high co-infection and reassortment rates. Swine housing density has been shown to correlate with H3 prevalence, which could enhance opportunities for coinfection and reassortment (Maes et al. 2000; Poljak et al. 2008). Lifespans of swine vary across production types, with sows living up to 2-3 years on average, and new susceptibles entering the population yearly, sustaining viral prevalence (Pitzer et al. 2016; Brown 2000; EFSA Panel on Animal Health and Welfare (AHAW) et al. 2022). Commercial swine are also moved frequently, enabling transmission between herds, resulting in a heterogenous immune landscape at the population level (Brown 2000). These patterns, in which susceptible hosts are concentrated, moved frequently, and periodically turn over, may increase the probability of reassortment, and of beneficial reassortants sustaining onward transmission (Thomas et al. 2024; Neveau et al. 2022; Marshall et al. 2013). Alternatively, it is also possible that the concentration of susceptible hosts could amplify the transmission of reassortant lineages, even in the absence of a fitness benefit, as observed in co-housed pigs where mutations persisted despite fitness costs (Murcia et al. 2012). Together, this combination of swine farming practices and swine immunity may facilitate the high reassortment and adaptive rates that we observe.

Our data recapitulate the frequent transmission of viral lineages between humans and swine, a long-existing phenomenon. We find enrichment of reassortment on human-to-swine host switches, which further supports the hypothesis that swine are more often recipients of human IAV lineages rather than sources (Nelson and Worobey 2018; Nelson and Vincent 2015; Neumann et al. 2009). While we cannot determine whether reassortment acts as the primary driver of the switch or a stabilizing force following the jump, its enrichment on human-to-swine switches suggests that reassortment is associated with host switches between these species, potentially facilitated by year-round IAV circulation in swine that is permissive to reassorting with human-adapted viruses. These findings suggest swine as unique progenitors of viral diversity, and underscore the importance of continued surveillance and risk assessment of swine IAV populations and at the human-swine interface.

This work reveals fundamental differences in how viral mixing is constrained between avian and swine hosts. In swine, NA and PB1 reassortments were significantly underrepresented relative to expectations under random reassortment, suggesting that swine reassortant viruses face HA-NA-PB1 balance constraints. This pattern matches the known evolutionary history of swine viruses, where North American swine H3N2 viruses received their HA, PB1, and NA segments from human H3N2 viruses in the late 1990s (Zhou et al. 1999). This human-derived constellation dates back to the 1968 pandemic strain which featured an HA-PB1 of avian origin paired with an NA from the H2N2 human lineage (Lindstrom et al. 2004). While H3N2 remains less prevalent in European swine, the H3N2 swine lineage possessed an HA-NA pairing from a human-adapted virus with internal segments donated from Eurasian avian H1N1 viruses (Zell et al. 2013; Castrucci et al. 1993). Phylogenetic analyses showing HA-NA diversity are linked in swine viruses further confirm this pattern (Zeller et al. 2021). These pairings may reflect an inherent functional constraint for circulation in mammalian hosts or epistatic interactions that co-evolved in these viruses following their introduction into the swine population (Nelson et al. 2014; Lowen 2017). In contrast, avian viruses showed an excess of NA reassortments, with novel NA subtypes introduced at rates exceeding those expected from observed viral circulation alone. Consistent with our findings that reassortment is largely neutral in avian hosts, this pattern may reflect incomplete surveillance of avian or environmental reservoirs. The NA subtype frequency calculations underlying our null model depend on observed sequence data, and substantial unsampled viral diversity could bias our estimates of expected reassortment probabilities. Understanding this dynamic will require more comprehensive field surveillance to better capture the true extent of viral diversity in natural avian reservoirs. Interestingly, there seems to be a lack of NA cross-reactivity in mallards, indicating complete sterilizing immunity against different NA subtypes is never established (Latorre-Margalef et al. 2013). This may make the reservoir host highly permissive to multi-subtype coinfection, which could directly facilitate the cocirculation of diverse NA subtypes.

The past four human influenza pandemics have emerged through reassortment events involving cross-species transmission, underscoring the importance of understanding host-specific reassortment and evolutionary dynamics (Taubenberger and Kash 2010). Here, we show that the H3Nx system provides a unique and underappreciated opportunity to observe how influenza viruses operate under different evolutionary pressures, highlighting its remarkable flexibility in evolutionary strategy depending on host type. These findings reveal that influenza evolution is fundamentally governed by host-specific ecology, epidemiology, and biology. Influenza relies primarily on reassortment in its avian reservoir but shifts towards accumulating adaptive substitutions upon endemic circulation in mammalian hosts, with the degree of this shift varying by host species. Our findings support swine as key hosts to surveil, particularly at the human-swine interface, where human-to-swine host switches frequently involve reassortment. Reassorted viruses in swine have a tendency to persist, which provides the opportunity to accrue mutations in the surface proteins that facilitate establishment in the new pigs, and drift from the original seeding human viruses (Vijaykrishna et al. 2011; Rajao et al. 2022). In avian hosts, reassortment instead generates and sustains extensive subtype diversity that co-circulates globally, where reassortant lineages are largely short-lived with limited fitness effects. However, this large and dynamic reservoir of circulating diversity provides repeated opportunities for mixing with strains with pandemic potential, including highly pathogenic influenza viruses (Damodaran et al. 2026). Overall, this work emphasizes the need to prioritize surveillance and biosecurity efforts, particularly in swine populations where reassortants may confer fitness benefits for mammalian adaptation and in avian populations where reassortment generates viral diversity capable of seeding future pandemic strains.

## Methods

### Data and phylogenetics

#### Initial H3Nx dataset curation and filtering

A comprehensive dataset of H3Nx genome sequences sampled from all non-human hosts, regions, and across time was downloaded from NCBI virus and GISAID (Khare et al. 2021) databases yielding 13,295 viral sequences sampled from birds, horses, camels, donkeys, dogs, seals, mink, swine, and cats collected between 1963 and 2024. Three human spillover H3N8 viruses were also included from GISAID (A/Guangdong/ZS-23SF005/2023, A/Henan/4-10/2022, A/Changsha/1000/2022).

To mitigate sampling biases across species and time periods, the following workflow using Snakemake v9.9.0 and Nextstrain v8.5.4 was followed. Sequences were filtered down to 5,023 full non-human genome based on these filtering parameters: (1) minimum sequence lengths per segment (HA: 1,600 bp; NA: 1,270 bp; NP: 1,400 bp; PA: 2,000 bp; PB1: 2,100 bp; PB2: 2,100 bp; MP: 900 bp; NS: 800 bp); (2) exclusion of sequences with collection dates prior to 1960; (3) removal of sequences with unknown country or region metadata; (4) removal of invalid characters. Sequences were grouped by year, country, host species, and subtype, and within each group the data were subsampled to 30 sequences per group. To ensure that all eight segments were represented by the same set of strains, subsampling was performed first for the HA segment, and then this filtering was applied to the other seven segments. Human seasonal H3N2 viruses collected between 1968-2023 were downloaded separately from NCBI (N=29,114) and filtered down to 1,078 sequences grouped by year and country. These sequences were then added to the H3Nx dataset to make global phylogenies, bringing the total filtered dataset size to 6,104 full genomes (5,023 non-human H3Nx genomes, 3 spillover human H3N8 genomes, 1078 human seasonal H3N2 genomes).

#### Phylogenetic reconstruction of H3Nx evolution

Filtered sequences for each segment were aligned to an H3N8 mallard sequence (A/blue-winged teal/Alberta/221/1978) using MAFFT v7.475. Maximum-likelihood phylogenetic divergence trees were inferred for each segment using IQ-TREE v2.4.0 with a GTR substitution model. Time-resolved trees were then constructed using TreeTime v0.11.4 using a coalescent time scale, where dates are inferred at internal nodes. Temporal outliers were identified using a clock filter (interquartile distance threshold: 4) and excluded. Nucleotide and amino acid sequences were reconstructed at internal nodes using TreeTime. Ancestral state reconstruction was performed for host type, bird order, country, region, and subtype via a discrete trait analysis. Segment trees are shown in Figure 1A and Supplementary Figure 18.

#### Host subtree construction

Currently circulating host groups (North American avian, Eurasian avian, swine, human H3N2 seasonal, equine, and canine H3N2 viruses) were parsed from the global H3Nx phylogeny based on their clustering within the HA phylogenetic tree. Here, we collapsed all geographic and phylogenetically distinct swine H3 lineages into a single “swine” clade. This approach enables a comparative analysis of viral evolution across mammalian and avian hosts at a global scale, but potentially obscures within-clade variability among swine lineages.

Each host-specific dataset was realigned using MAFFT with host-specific references for each segment. Maximum-likelihood phylogenetic divergence trees were inferred for each segment using IQtree with a GTR substitution model. Clock rates were estimated using TreeTime’s clock model function in python. These alignments and trees were used for the subsequent mutation and reassortment analyses.

### Adaptive substitution rates

#### Modified McDonald-Krietmand test (Bhatt method)

For each host group, we performed a modified McDonald-Krietman (MK) test on the HA, NA, and PB1 genes(Kistler and Bedford 2023; Bhatt et al. 2011). Here, PB1 was chosen as a conserved reference gene, since as a core polymerase subunit under strong purifying selection and not directly targeted by host immunity, it is not expected to accumulate adaptive substitutions to the same degree as HA and NA (Kistler and Bedford 2023; Bhatt et al. 2011). This test calculates adaptive substitution rates by measuring the accumulation of nonneutral substitutions above a neutral expectation and was specifically developed for temporally-sampled viral genomes. Aligned viral sequences were binned into overlapping time windows of at least 3 years, with a minimum of 3 sequences per time window. The test uses an updating outgroup, which is first taken as the consensus of sequences in the earliest time window. This outgroup was updated for future time windows when fixations at each site occur, to account for recurring mutations. This outgroup is then compared to the aligned sequences in each subsequent time window (the ingroup). Across each site, genetic differences between the outgroup and ingroup are noted, and the number of adaptive substitutions is calculated as the sum of nonsynonymous fixations and near fixations that exceed the neutral expectation. Reflecting influenza’s high mutation rate, the neutral expectation is measured by counting: 1) all mid-frequency (15-75%) polymorphisms, 2) synonymous near fixations (>75%), and 3) synonymous fixations. Adaptive substitutions per codon are then plotted as a point estimate for each time window, where the slope of the linear regression fitting these estimates is calculated as the rate of adaptation (Supplementary Figure 3, Figure 1B). Adaptive substitutions rates are measured as adaptive substitutions per codon per year × 10^−3^.

The alignments for the host-specific subtrees were further filtered to include only competing viruses, defined as viruses co-circulating within the same host and geographic region. For avian viruses, competing viruses for the NA segment were additionally required to belong to the same subtype. This filtering strategy accounts for this test’s sensitivity to population structure and ensures that adaptive substitution rates reflect selection pressures within epidemiologically relevant viral populations. Specifically, the swine host group was subdivided into separate European and North American populations and North American and Eurasian avian NA lineages were stratified by subtype (N2 and N8). These population-level groupings merge independent introductions across these regions to enable larger scale comparisons of viral evolution among these hosts. Reassortant strains identified through TreeSort were excluded to prevent sequence differences arising from reassortment events from biasing selection estimates.

Code for this test was adapted from github.com/blab/adaptive-evolution/blob/master/adaptive-evolution-analysis.

### TreeSort pipeline and reassortment rates

#### Overview of TreeSort and TreeSort Pipeline

TreeSort implements a phylogenetic incongruence approach to identify reassortment by designating a single segment as a reference and systematically comparing the evolutionary histories of the remaining segments against it. In the absence of reassortment, these phylogenies are expected to be largely congruent where deviations from this expectation are interpreted as evidence of reassortment. To do this, TreeSort estimates segment-specific molecular clock rates and uses these to define probability thresholds for detecting reassortment events along branches of the reference phylogeny. This method produces a single annotated tree in which inferred reassortment events and the reassorting segments are mapped. Here, a reassortment event is defined as 1 or more segments reassorted relative to HA on a single branch.

Although TreeSort is highly informative, a single run yields only one annotated phylogeny, thereby limiting inference to point estimates for different parameters (e.g. reassortment rates). To address this limitation, we developed a pipeline to run TreeSort in replicate (available at github/moncla-lab/treesort-pipeline). This approach allows for more robust reassortment inference by incorporating statistical uncertainty in tree building and TreeSort’s respective probability estimations.

The pipeline was generated using a Snakemake workflow and proceeds as follows: TreeSort is run once to generate a binarized backbone tree annotated with reassortment events, which serves as the fixed reference phylogeny for all subsequent replicate runs. In each replicate, new divergence trees are generated for the challenge (non-reference) segments while the *--no-collapse* flag is applied to ensure all TreeSort outputs retain identical topology to the backbone tree. For these analyses, 1000 replicates were performed. Reassortment rates (reassortments per lineage per year) are calculated for each replicate and compiled in a summary log file.

After running TreeSort in replicate, a summary JSON file is generated that records the reassortment support value for each node, defined as the proportion of runs in which a reassortment event was inferred. Support values are calculated independently for each segment, such that a node involved in a multi-segment reassortment event (e.g., PB2, PB1, and PA jointly) may have differing support values across the reassorting segments. A summary reassortment tree is also generated, annotated only with high-support reassortments (reassortments inferred in ≥95% of the runs). The resulting summary reassortment tree is able to be visualized via the Nextstrain Auspice interface using the cladeset-mapping tool (described in github/moncla-lab/treesort-cladeset-mapping), enabling interactive exploration of reassortment event inferences and their support across the reference phylogeny.

When TreeSort cannot confidently assign a reassortment event to a specific branch, it labels both child branches with an uncertain tag (e.g., ?PB2 to indicate ambiguity regarding the PB2 segment, see https://github.com/flu-crew/TreeSort for details). In such cases, one of the two child branches was called at random to be reassorted.

Custom Python scripts (v3.12.11) are used throughout the pipeline: (1) converting TreeSort replicate outputs into Newick trees, (2) calculating reassortment rates, (3) recording individual reassortment events and resolving uncertain calls across replicates into individual JSON files, (4) aggregating results from individual JSONs into a summary file, and (5) mapping consensus reassortments back to a nonbinarized tree for Nextstrain visualization.

#### Reassortment rates and North American avian subsampling

As part of the TreeSort pipeline, the reassortment rate is calculated at each replicate as the total number of reassortments divided by total tree length, and then scaled by the evolutionary rate of the reference segment (see (Markin et al. 2025) for details). For each host group, we calculated the mean rate and standard deviation across replicates (Figure 2A). Reassortment rates are in units of reassortments per year per lineage, enabling direct comparison across host groups.

To assess the sensitivity of reassortment inferences to dataset size, we generated serially subsampled datasets of North American avian H3Nx viruses, ranging from N=400 to N=1,100 sequences in 100-sequence increments. Sequences were subsampled by year and region. Regions were defined as sequences sampled from Africa, Europe, North America, China, South Asia, Japan/Korea, Oceania, South America, and West Asia. For each dataset size, 5 independent subsampling trials were performed, and reassortment rates were calculated using the full TreeSort replicate pipeline described above (Supplementary Figure 6).

#### Null distribution - shuffled reassortment summary tree

To assess whether observed reassortment dynamics were ever consistent with reassortment under a random model, we generated a null distribution of reassortment for each host group and the global H3Nx datasets. For each host, we randomly redistributed all high-support reassortment events uniformly across the branches of the tree, such that each branch had an equal probability of being assigned a reassortment event regardless of branch length or tree structure. This procedure was repeated 1000 times for each group, generating a distribution of 1000 trees in which the number of reassortment events and tree topology were conserved, but placement was randomized.

#### Flyway analysis for North American avian viruses

To determine whether reassortment events occurred more frequently in specific flyways than expected by chance, we performed a discrete trait analysis on the North American avian dataset to infer flyway annotations at internal nodes. Flyway assignments for terminal taxa were based on sampling location according to US Fish and Wildlife Service Administrative Flyway classifications (U.S. Fish & Wildlife Service 2023). We then calculated the frequency of each flyway among all sequences as a proportion of the total, which served as the null expectation probability for each flyway.

For each flyway, we counted the number of reassorted nodes assigned to that flyway as a proportion of the total number of reassorted nodes. We then used a two-sided binomial test to determine whether the observed frequency of reassortment events in each flyway differed significantly from the expected frequency based on the overall strain distribution. Multiple hypothesis testing was corrected using the Bonferroni method across all flyways, with significance threshold set at α = 0.05. We repeated this flyway analysis on the null distribution of trees described earlier.

### Segment reassortment dynamics

#### Null distribution for segment-specific reassortments

To assess whether gene segments reassort nonrandomly, we generated a null model based on the observed reassortment events. The distribution of event sizes (the number of segments involved per reassortment event) was first quantified from the observed data. Simulations were then performed in which the observed distribution of event sizes was maintained, while the identities of reassorting segments were randomly assigned by sampling without replacement. This procedure was repeated 1,000 times to generate a null distribution of randomly reassorting segments. Observed reassortment frequencies were then compared to the simulated null distributions using empirical two-sided p-values, accounting for small sample size using a Monte Carlo correction and multiple testing using Bonferroni method (α = 0.05). P-values were calculated as *p = (r + 1) / (n + 1)*, where *r* represents the count of simulated values with absolute deviation from the mean as extreme or more extreme than the observed value in either direction, and n equals the total number of simulations (n = 1,000).

#### Calculating within- and between-reassortment probabilities for avian host groups

To determine whether observed within- and between-subtype reassortment counts deviated from expectations based on subtype circulation, expected reassortment probabilities were calculated. For both North American and Eurasian avian datasets separately, the frequency of each NA subtype was determined by counting the number of strains belonging to each subtype and converting these counts into proportions of the total datasets. Expected within-subtype reassortment probabilities were estimated by summing the squared frequencies of each NA subtype, corresponding to the probability of randomly selecting two viruses belonging to the same subtype. Expected between-subtype probabilities were calculated as the complement of the within-subtype probability (1-P(within)), representing the probability of selecting two viruses of different subtypes. In North American avian viruses, the expected within-subtype reassortment probability is 38%, and the expected for between-subtype is 62%. For Eurasian viruses, the expected within-subtype reassortment probability is 41%, and the expected for between-subtype is 59% (Supplementary Figure 10). These probabilities were compared to observed counts under a binomial model.

#### Calculating within- and between-subtype divergence thresholds

To distinguish within- and between-subtype NA reassortment events, pairwise nucleotide divergence was calculated among all NA sequences within the North American and Eurasian avian datasets. Low-quality sequences (those containing fewer than 80% valid nucleotide sites) were filtered out. Pairwise divergence was then calculated for all possible sequence pairs within each dataset, labeled within-subtype when both sequences belonged to the same NA subtype and between-subtype when sequences belonged to different NA subtypes. Within- and between-subtype divergence thresholds were determined from these empirical distributions of pairwise NA sequences divergences. The within-subtype divergence threshold was defined as the maximum pairwise divergence observed among sequences belonging to the same NA subtype, whereas the between-subtype threshold was defined as the minimum pairwise divergence observed among sequences belonging to different NA subtypes. The within-subtype threshold was used as the upper bound for classifying NA reassortments as within-subtype events.

#### Classification of NA reassortments as within- or between-subtype events

TreeSort calculates how diverged a reassorting segment is relative to the parental segments, measured as the number of nucleotide differences. For example, an annotation of PB2(136) denotes acquisition of a PB2 segment that differed by 136 nucleotides from the parental PB2 segment (see https://github.com/flu-crew/TreeSort for details). Using only high-support NA reassortments, the inferred NA divergence values inferred by TreeSort were compared to the divergence thresholds calculated above. Reassortment events with TreeSort-inferred NA divergence values less than or equal to the within-subtype upper bound were classified as within-subtype reassortments. Reassortment events exceeding this bound were classified as between-subtype reassortments. Among 173 NA reassortment events identified in North American avian viruses, 35 involved reassortment of NAs from the same subtype, while 138 involved introduction of NA segments from different subtypes. Among 148 NA reassortment events identified in Eurasian avian viruses, 39 involved reassortment of NAs from the same subtype, while 109 involved introduction of NA segments from different subtypes.

#### Testing if novel NAs are brought in more than expected by chance

To test whether within- and between-subtype NA reassortment events deviated from expectations under a model of random reassortment, observed counts were tested under a binomial model. The expected within- and between-subtype reassortment probabilities derived from circulation frequencies were used as success probabilities. The observed number of “successes” was taken as the number of within- or between-subtype reassortments classified above. The binomial distributions were then evaluated over all possible outcomes to generate the expected probability mass function for within- and between-subtype reassortments under a null model. The p-value was computed as the cumulative probability of observing the actual number of within- and between-subtype events or fewer under the null model. The binom function from the scipy.stats library was used for these tests.

#### Binomial sensitivity analysis

To evaluate whether potential detection bias against more highly divergent NA reassortment events could affect inference of within- and between-subtype reassortment, we performed a binomial model sensitivity analysis. First, we estimated the proportion of within-subtype NA sequence pairs that fell below the minimum divergence observed among NA reassortment events inferred by TreeSort. This minimum divergence threshold was taken as an empirical lower bound for detectable reassortment events, and the fraction of within-subtype sequences pairs below this threshold was used as an estimate of potential underdetection.

To assess the sensitivity of our findings to this underdetection, we then systematically increased fractions of observed within-subtype while keeping the total number of events constant. Across a range of scenarios (1-12% additional within-subtype events), binomial tests were recomputed using the same null probabilities derived from cocirculation. For each scenario, we evaluated whether the observed excess of between-subtype remained statistically significant as within-subtype events increased. We identified the maximum proportion of within-subtype underdetection that could be tolerated before statistical significance was lost (α = 0.05). Here, the significance of our findings was robust up to 11.6% underdetection in North American and 7.4% in Eurasian avian viruses (Supplementary Figure 13).

### Fitness effects of reassortment

#### Reassorted persistence analysis

To assess whether reassortment confers a fitness advantage within avian and swine host groups, we quantified reassortant lineage persistence as proxy for viral fitness. For each high-support reassorted internal node inferred by the TreeSort pipeline, we calculated the maximum distance to either the next downstream reassortment event or a terminal leaf node. Branch lengths were scaled to time in years by dividing by the host-specific molecular clock rate.

We compared persistence times of reassorted lineages to those for nonreassortant lineages. Nonreassortant lineage persistence was measured as the maximum distance from a non-reassorted node to the next terminal leaf or reassortment event. Note that this calculation does not account for whether the nonreassortant lineage descends from a recent reassortment event. Persistence times were binned into annual intervals to create discrete persistence categories (0–1 years, 1–2 years, etc.). For each host group and lineage type (reassorted vs. non-reassorted), we calculated the proportion of lineages falling into each persistence bin relative to the total number of lineages (Supplementary Figure 16). A similar analysis was performed to test whether observed reassortant persistences differed from random expectation. Using the distribution of null reassortment trees described earlier, we calculated mean null reassortant lineage persistences and the 95% confidence intervals for each persistence bin.

#### Long-term fitness effects of reassortment

To determine whether reassortment is associated with the long-term survival of viral lineages, we assessed the likelihood that reassorted nodes produce descendants surviving *x* years into the future (*x* = 0.5, 2, 6, and 10 years post-reassortment). We only evaluated internal nodes with an estimated age at least equal to the specified time interval. For each qualifying node, we calculated the maximum temporal distance from the node to its most recent descendant leaf. Nodes were classified as “fit” if they produced descendants reaching or extending beyond the specified time threshold, or “unfit” if all descendants died out before the threshold.

For each time interval and host group (Swine, Eurasian, and North American avian), a 2×2 contingency table was constructed crossing lineage type (reassorted vs non-reassorted) and survival outcome (fit vs unfit). We performed a two-sided Fisher’s exact test using the *fishers_exact* function from the *scipy.stats* library. Odds ratio and 95% confidence intervals were calculated from each contingency table confidence intervals were computed using the log-transformed odds ratio function from the *math* library.

We also performed these Fisher’s exact tests on the distribution of null, shuffled reassortment trees described earlier. For each host group and time interval, we identified how often null trees yielded statistically significant results (α = 0.05) and compared the distribution of odds ratios across the null trees to the actual odds ratios (Supplementary Figure 17).

### Host-switching analysis

#### Reassortment enrichment on host-switched branches

We identified host switches in the global H3Nx phylogenetic tree by comparing host assignments between parent and child nodes using inferences from the discrete trait analysis. If a parent host assignment differed from its child node, then this was considered a host-switched branch. For each internal node, we extracted its reassortment status and classified all host switches into four categories: reassorted host switches, non-reassorted host switches, reassorted non-switches, and non-reassorted non-switches. To test whether reassortment events were statistically enriched on host-switching branches, we constructed a 2×2 contingency table and performed a two-sided Fisher’s exact test. Odds ratios were computed using the *fishers_exact* function from the *scipy.stats* library and 95% confidence intervals were calculated using the log-transformed odds ratio function from the *math* library. Here, an odds ratio > 1 and p-val < 0.05 indicates reassortment enrichment on host-switched branches. This analysis was also conducted separately for the North American and Eurasian avian datasets but for transitions between avian orders (e.g. galliform to anseriform) where no significance was found.

To determine whether the observed association between reassortment and host-switching varied by transition type, we compared observed reassorted transitions in the global H3Nx tree to transitions in the distribution of null trees described earlier. For each null tree, we identified all host switches occurring on reassorted internal nodes and stratified them into 3 categories: avian-to-mammal, mammal-to-mammal, and mammal-to-avian transitions. We aggregated counts across the 1,000 null trees to generate null distribution for each host transition. Empirical p-values were calculated as *p = (r + 1) / (n + 1)* where *r* represents the number of null trees with a host transition count as extreme or more extreme than the observed count and *n* equals the total number of null trees (n=1000) (Figure 5B).

### Code availability and reproducibility

All code developed and used in this project are available in the github.com/moncla-lab/h3nx-paper git repo. The TreeSort pipeline is available with documentation and example data at github.com/moncla-lab/treesort-pipeline. A public, interactive version of the H3Nx tree is available from the Moncla lab Nextstrain groups page at: nextstrain.org/groups/moncla-lab/h3nx/ha. All data that were used in this analysis were sourced from public databases. The acknowledgement table for GISAID isolates used in this analysis is provided in Supplementary Table 2, which can also be found at GitHub (https://github.com/moncla-lab/h3nx-paper).

## Acknowledgments

We gratefully acknowledge all data contributors, i.e., the Authors and their Originating laboratories responsible for obtaining the specimens, and their Submitting laboratories for generating the genetic sequence and metadata and sharing via the GISAID Initiative, on which this research is based.

Support for this project was provided by the Pew Charitable Trusts, the Margaret Q. Landenberger Research Foundation, the National Institute of Allergy and Infectious Diseases, National Institutes of Health, Department of Health and Human Services (contract numbers 75N93021C00015 and 75N93021C00016), and the United States Department of Agriculture, Agricultural Research Service (ARS project number 5030-32000-231-000D). USDA is an equal opportunity provider and employer.

