## supplemental figure 1-18, supplemental table 1 for "Host type governs influenza evolutionary strategy across reservoir and spillover hosts"

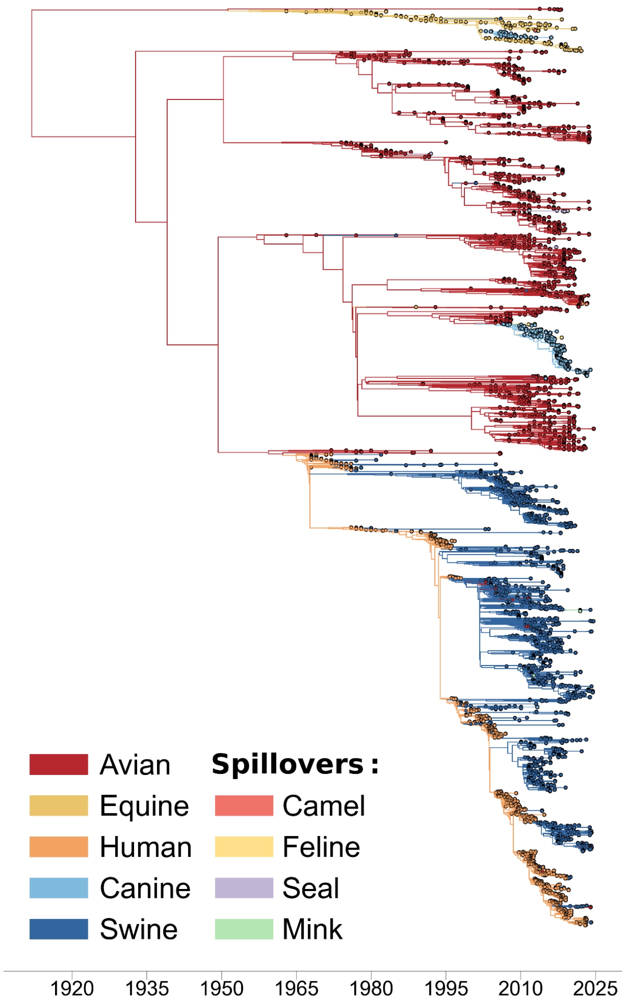


**Supplemental Figure 1: H3Nx viruses have jumped hosts multiple times.** An enlarged view of the HA phylogeny (N=6,130) colored by host type, where each leaf is a unique strain and each internal node is an inferred ancestor. Notable host switches are highlighted. The 95% confidence interval for the TMRCA estimates are between 1910-05-22 – 1917-09-14.

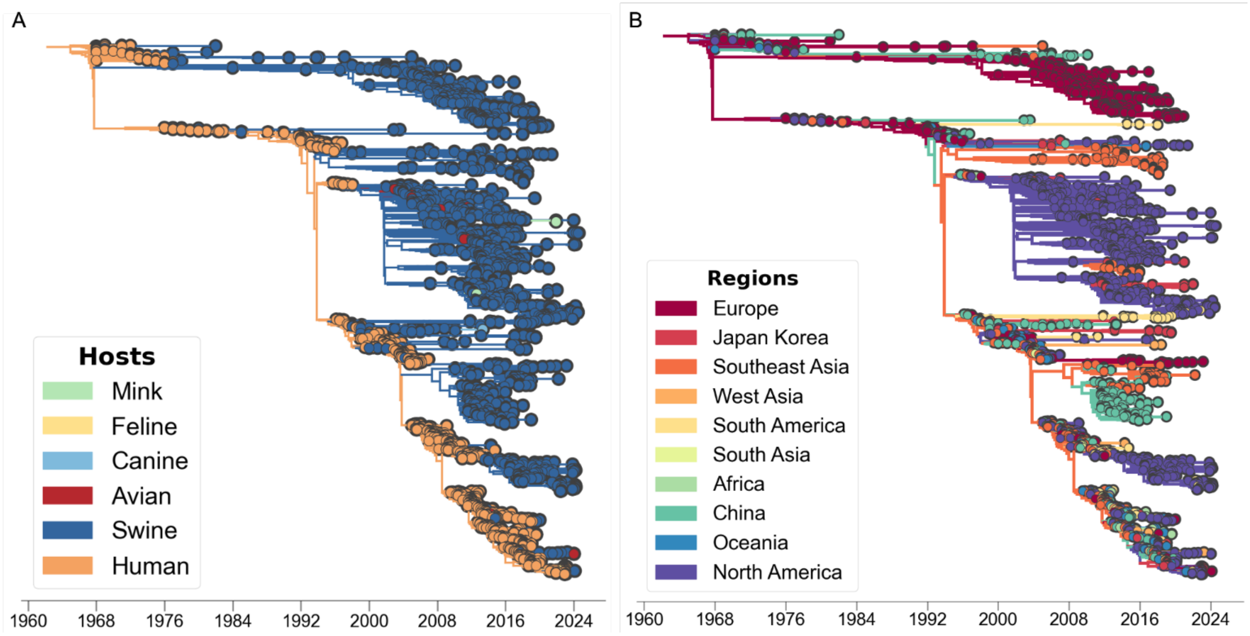


**Supplemental Figure 2: There have been repeated spillbacks between humans and swine.** ***A)*** The HA phylogeny subsetted to viruses circulating in human and swine populations (N=3,140) colored by host type, where each leaf is a unique strain and each internal node is an inferred ancestor. The 95% confidence interval for the TMRCA estimates are between 1964-03-15 – 1965-06-28. ***B)*** The same tree is colored by region.

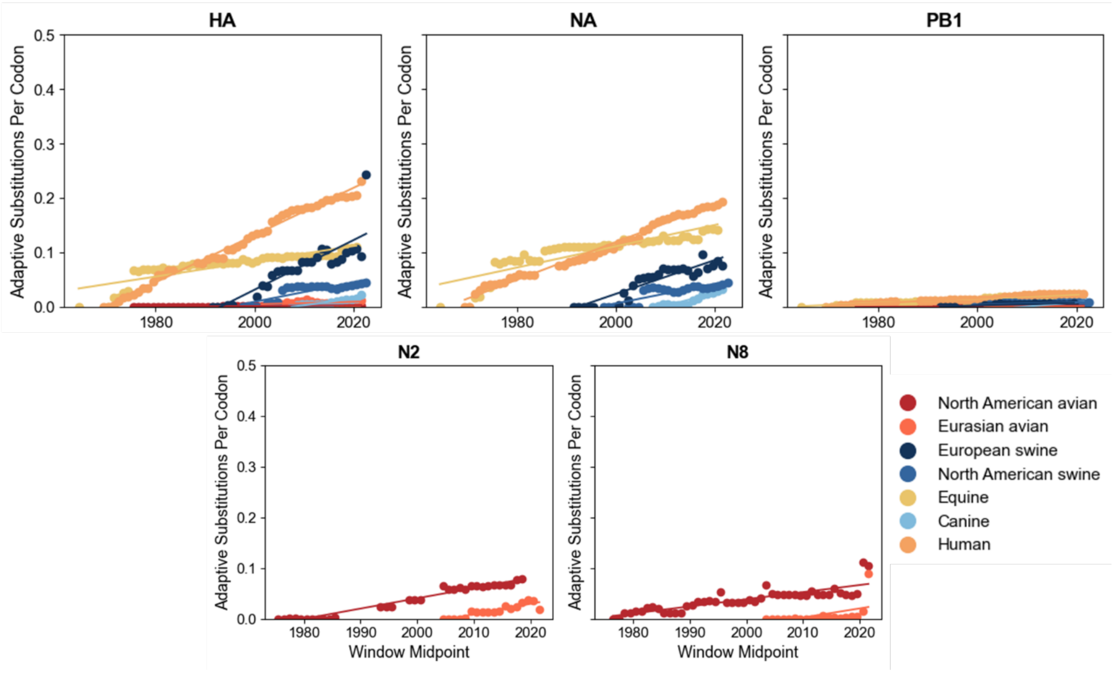


**Supplementary Figure 3: Estimated accumulation of adaptive mutations per codon across sliding time windows for HA, NA, and PB1 gene segments.** Each point represents the estimated adaptive substitution rate within a given time window, calculated using a modified McDonald-Kreitman test designed for temporally-sampled viral genomes. Lines show linear regressions fitted per host group, with the slope representing the rate of adaptive substitution accumulation over time. NA is split into N2 and N8 lineages for North American and Eurasian avian viruses. Data is colored by host type.

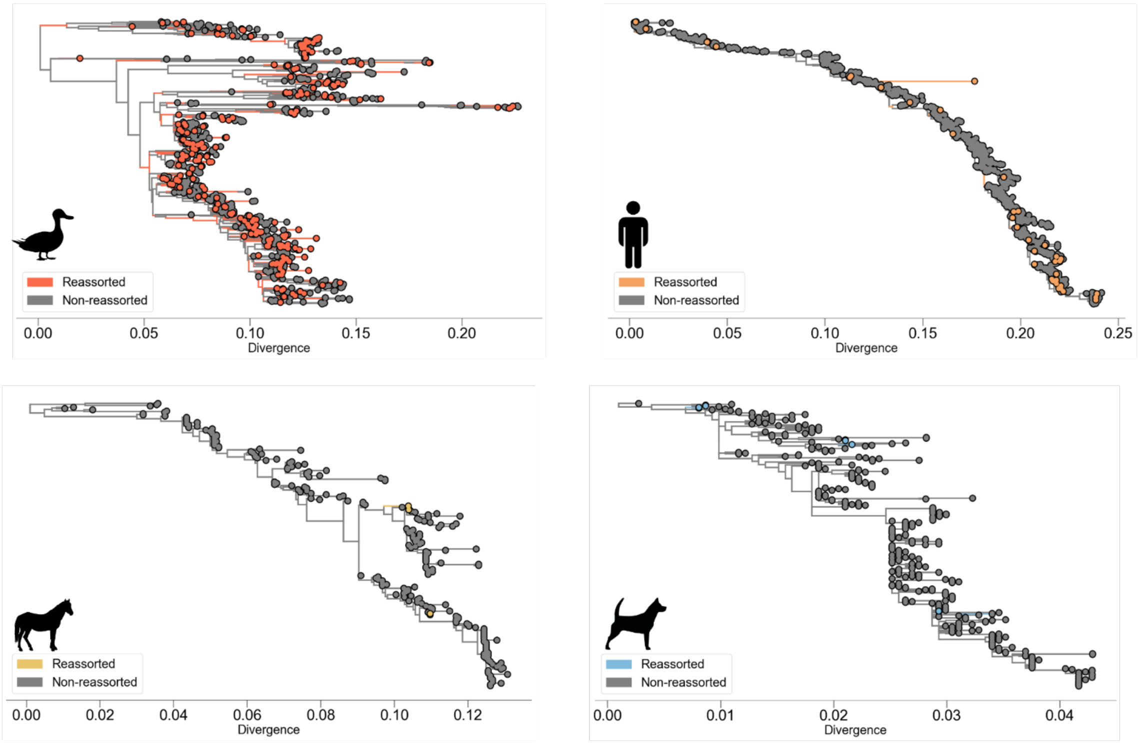


**Supplementary Figure 4: Reassortment vary across hosts.** Summary TreeSort trees shown for Eurasian avian, human, equine, and canine host groups. Branches are colored to indicate reassortment events supported at >= 95%.

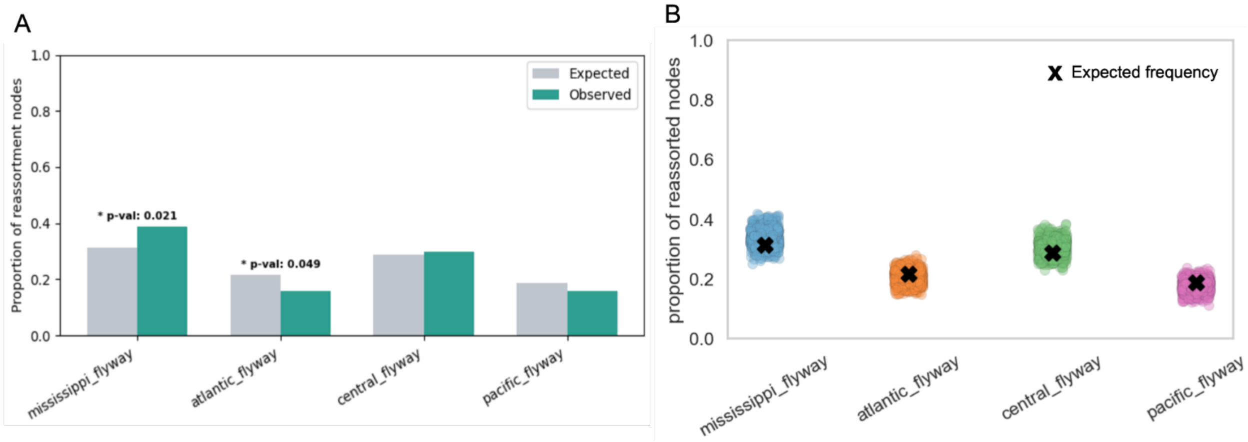


**Supplementary Figure 5: Reassortment events show non-random geographic distribution across North American flyways.** ***A)*** Observed versus expected proportion of reassorted nodes in each North American avian flyway, determined by binomial tests with expected proportions derived from flyway frequencies at tree tips. Asterisks indicate significance after Bonferroni correction. These results show a modest enrichment of reassortment in the Mississippi flyway (p-val: 0.021) but a slight depletion of reassortment in the Atlantic flyway (p-val: 0.049). ***B)*** Distribution of flyway proportions among reassorted nodes across 1,000 null trees generated by randomly shuffling reassortment event positions while preserving tree topology. Each point represents one replicate, colored by flyway. The black X indicates the observed proportion of reassorted nodes in each flyway from the observed data. No flyway showed significant deviation from the null distribution in the permutation analysis, suggesting observed results are not artifacts of sampling.

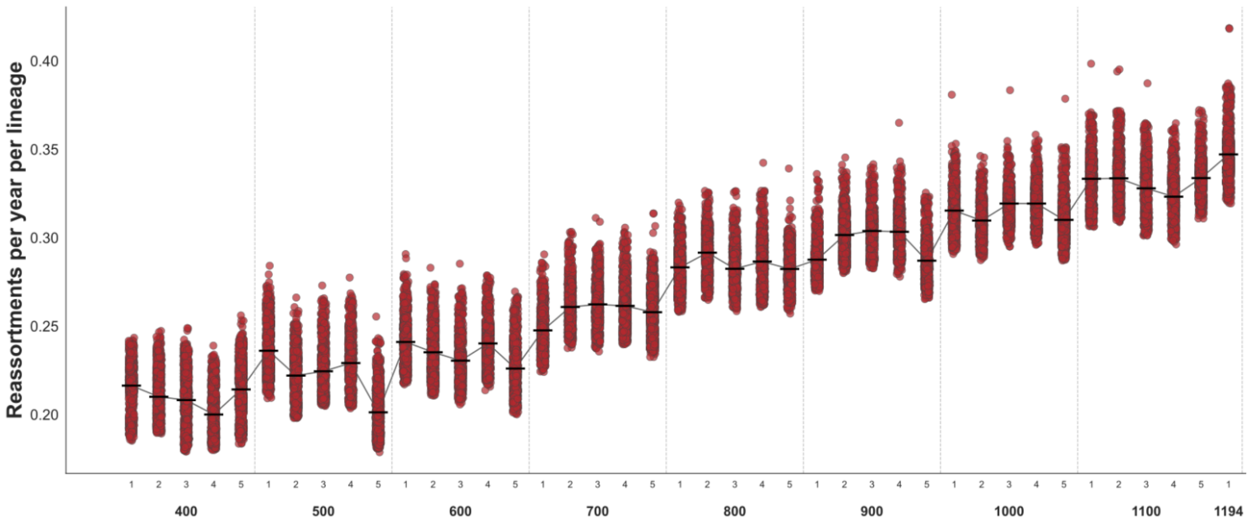


**Supplemental Figure 6: Reassortment rates for North American avian viruses increase with dataset size, but still exhibit the highest rates among host groups.** Reassortment rates (reassortments per year per lineage) were estimated from serially subsampled datasets of North American avian viruses ranging from N=400 to N=1100 sequences (100-sequence increments, 5 trials each), plus the full dataset (N=1194). Each point represents a reassortment rate calculated for an individual TreeSort replicate. All 1000 replicates for each subsampling and trial are shown, with the mean values indicated with a horizontal line. The line connecting means illustrates the positive relationship between dataset size and inferred reassortment rate, suggesting absolute rates are sensitive to sampling depth. Despite this, the relative ranking of reassortment rates among host lineages was consistent across all dataset sizes, supporting the use of relative comparisons among hosts even when absolute rates may vary.

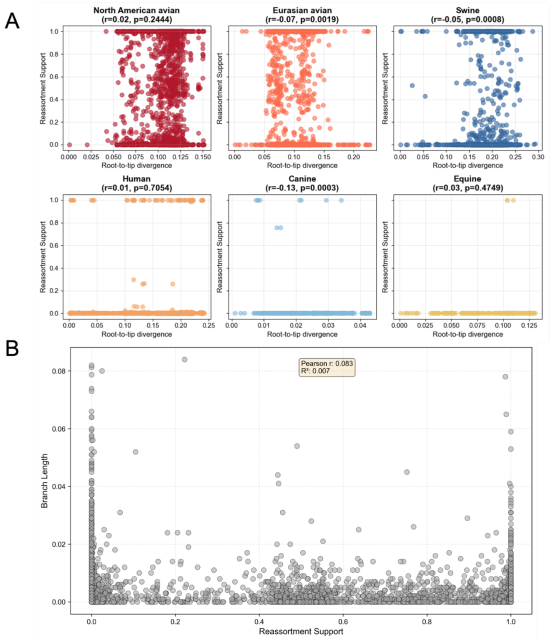


**Supplemental Figure 7: Reassortment inferences are robust to divergence.** For each detected reassortment event, support scores (proportion of TreeSort replicates calling the event) are plotted against the **A)** root-to-tip divergence of the corresponding node for each host group. Pearson correlation coefficients and p-values are indicated for each clade. No significant correlation was observed for North American avian ($r = 0.02$, $p = 0.24$), human ($r = 0.01$, $p = 0.71$), or equine ($r = 0.03$, $p = 0.47$) groups. Small but statistically significant negative correlations were detected in Eurasian avian ($r = -0.07$, $p = 0.002$), swine ($r = -0.05$, $p = 0.0008$), and canine ($r = -0.13$, $p = 0.0003$) groups. However, the effect sizes (r2) are small across all clades, and we do not interpret these as evidence of systematic bias in TreeSort's detection of reassortment events. **(B)** For the global H3Nx tree, reassortment support showed minimal correlation with branch length ($r = 0.083$, r2 = 0.007), indicating that reassortment detection is independent of branch divergence.

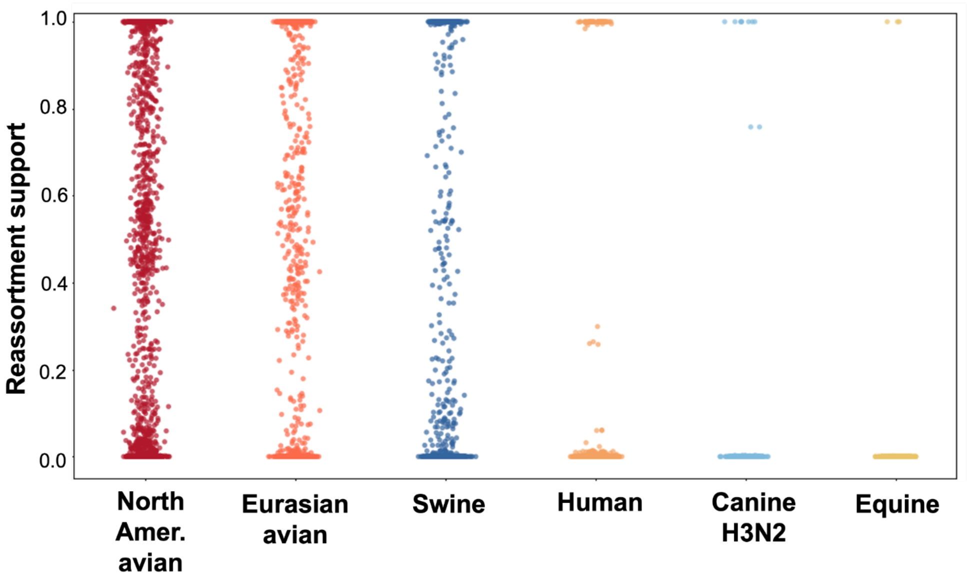


**Supplemental Figure 8: Reassortment support values per node vary across TreeSort replicates in highly reassorting host groups.** For each detected reassortment event, support values reflect the proportion of TreeSort replicates in which that event was called. Each point represents a leaf or node; values range from 0 (never called) to 1 (called in all replicates). North American avian, Eurasian avian, and swine groups show greater spread in support values compared to the human, canine, equine hosts. Reassortments called consistently across replicates are more likely to represent true reassortment signal, while low-support events may reflect spurious inferences called in individual replicates.

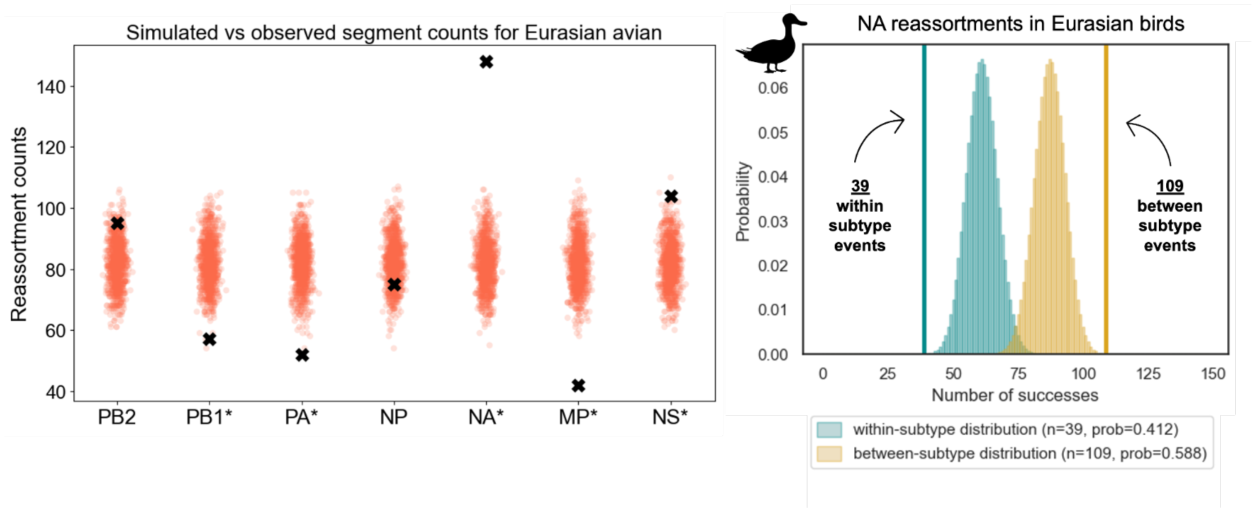


**Supplementary Figure 9: Eurasian avian segments exhibit host-specific compatibility with HA.** ***A)*** Segments reassort nonrandomly relative to a null model in Eurasian avian viruses. Data points represent simulated reassortment counts for each segment under random sampling while observed counts are denoted with a black X. Asterisks indicate p-value < 0.05 after Bonferroni correction. ***B)*** Binomial testing revealed a significant excess of between-subtype NA reassortments (p-value = 0.0001). Success probabilities for the binomial model were defined as the within-subtype (0.412) and between-subtype (0.588) NA reassortment probabilities. Observed within-subtype (blue, 39 events) and between-subtype (yellow, 109 events) NA reassortments are indicated by vertical lines.

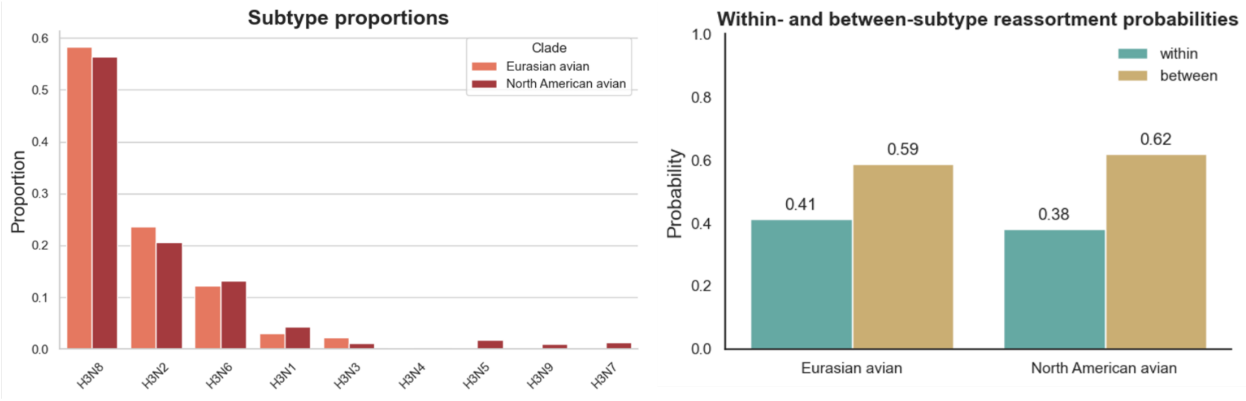


**Supplemental Figure 10. Expected within- and between-subtype reassortment probabilities based on NA subtype cocirculation. (A)** Proportions of NA subtypes circulating in North American avian and Eurasian avian virus populations. Subtype frequencies were calculated by counting strains belonging to each NA subtype and converting to proportions of the total. **(B)** Expected probabilities of within-subtype and between-subtype reassortment events under a null expectation of random reassortment. Expected within-subtype reassortment probability was calculated as the sum of squared subtype frequencies (reflecting the probability of independently sampling two strains of the same subtype), while expected between-subtype probability was calculated as 1 minus the within-subtype probability. In North American avian viruses, the expected within-subtype reassortment probability is 38%, and the expected for between-subtype is 62%. For Eurasian viruses, the expected within-subtype reassortment probability is 41%, and the expected for between-subtype is 59%.

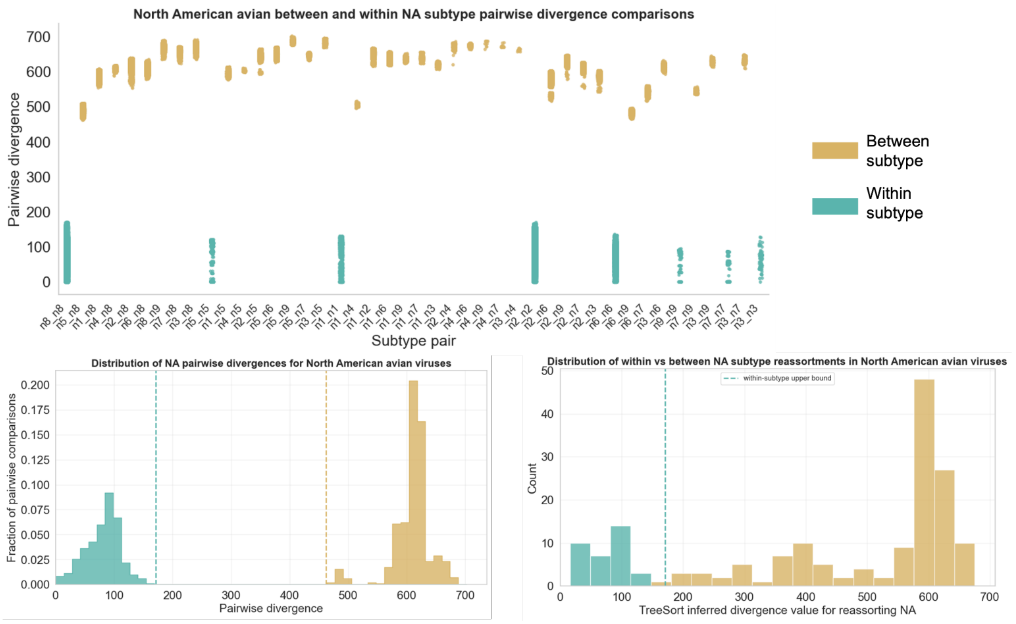


**Supplemental Figure 11. Classification of within- versus between-subtype NA reassortment events in North American avian viruses. (A)** Pairwise divergence among all NA sequences in North American avian viruses, stratified by subtype pairing (within-subtype pairs in blue, between-subtype pairs in yellow). Clear separation between the two distributions allows definition of a divergence threshold distinguishing within- and between-subtype comparisons. **(B)** Distribution of pairwise divergences, showing the threshold (dashed lines) that separates within-subtype (blue) from between-subtype (yellow) comparisons. **(C)** Distribution of TreeSort-inferred divergence values for reassorting NA segments, classified as within-subtype (≤ threshold, blue) or between-subtype (> threshold, yellow) based on the cutoff defined in panel B. Among 173 NA reassortment events identified in North American avian viruses, 35 involved reassortment of NAs from the same subtype, while 138 involved introduction of NA segments from different subtypes.

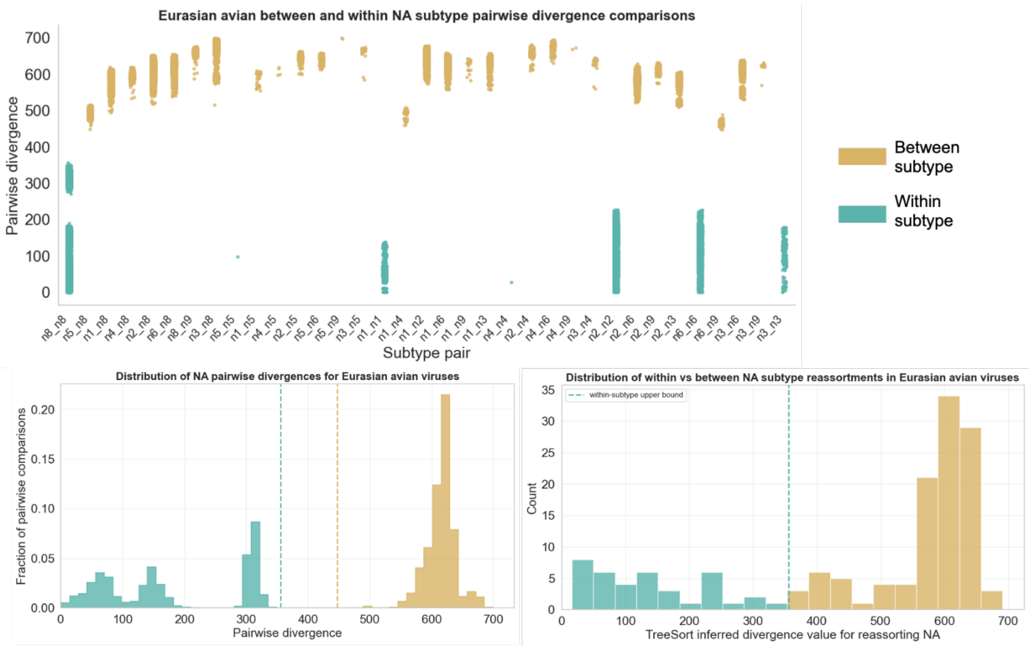


**Supplemental Figure 12. Classification of within- versus between-subtype NA reassortment events in Eurasian avian viruses. (A)** Pairwise divergence among all NA sequences in Eurasian avian viruses, stratified by subtype pairing (within-subtype pairs in blue, between-subtype pairs in yellow). Clear separation between the two distributions allows definition of a divergence threshold distinguishing within- and between-subtype comparisons. **(B)** Distribution of pairwise divergences, showing the threshold (dashed lines) that separates within-subtype (blue) from between-subtype (yellow) comparisons. **(C)** Distribution of TreeSort-inferred divergence values for reassorting NA segments, classified as within-subtype (≤ threshold, blue) or between-subtype (> threshold, yellow) based on the cutoff defined in panel B. Among 148 NA reassortment events identified in Eurasian avian viruses, 39 involved reassortment of NAs from the same subtype, while 109 involved introduction of NA segments from different subtypes.

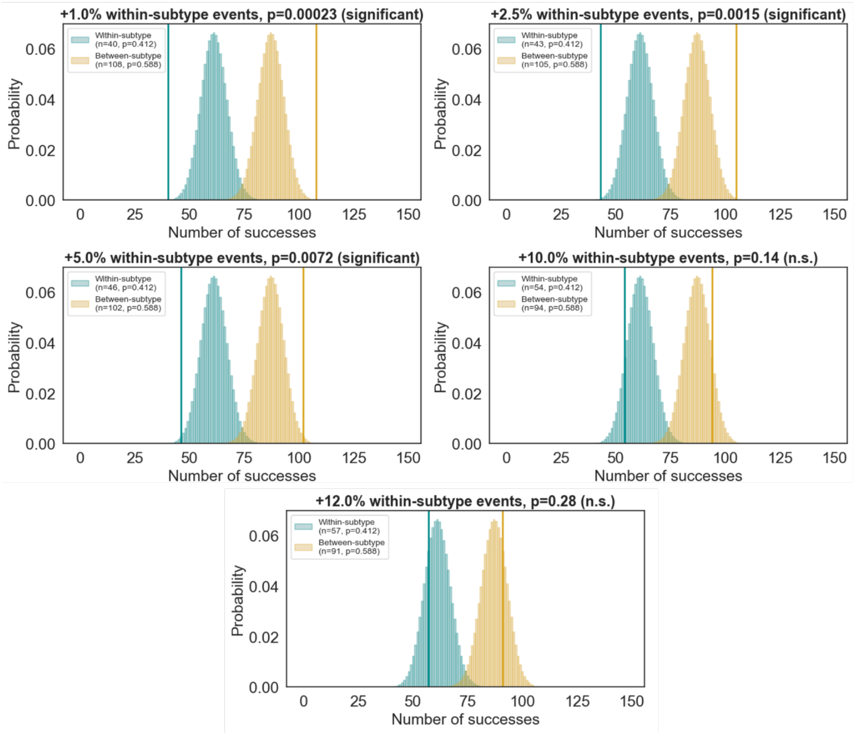


**Supplementary Figure 13. Binomial sensitivity analysis for Eurasian avian viruses.** Each panel shows the binomial null distributions for within-subtype (blue) and between-subtype (yellow) reassortment counts after increasing within-subtype events (1–12%), simulating the potential underdetection of these events by TreeSort. Vertical lines indicate the adjusted observed counts at each correction level. Between-subtype remains significant up to a underdetection threshold of 7.4%. As the estimated underdetection limit from pairwise divergence comparisons is 1.6%, the observed enrichment of between-subtype reassortments is considered robust to detection bias.

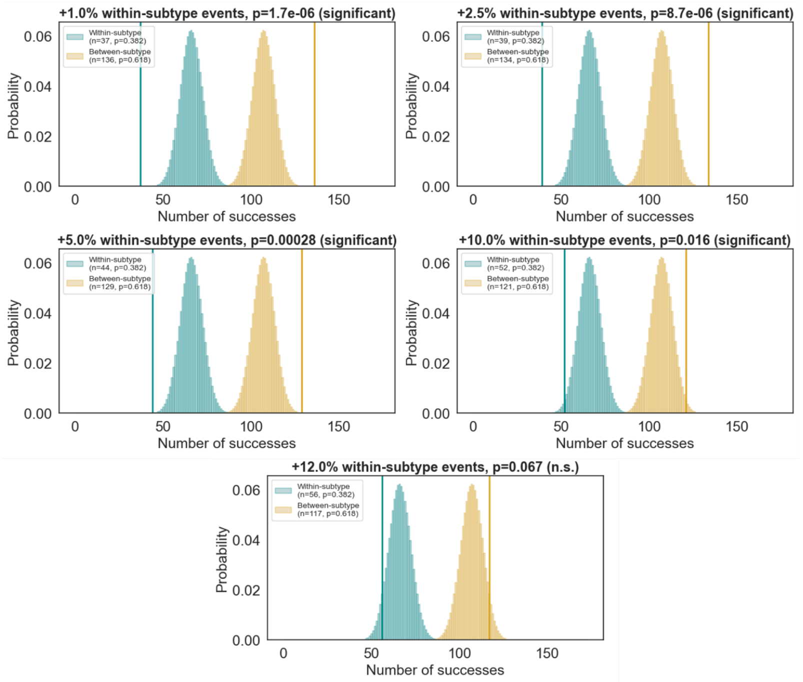


**Supplementary Figure 14. Binomial sensitivity analysis for North American avian viruses.** Each panel shows the binomial null distributions for within-subtype (blue) and between-subtype (yellow) reassortment counts after increasing within-subtype events (1–12%), simulating the potential underdetection of these events by TreeSort. Vertical lines indicate the adjusted observed counts at each correction level. Between-subtype remains significant up to a underdetection threshold of 11.6%. As the estimated underdetection limit from pairwise divergence comparisons is 2.4%, the observed enrichment of between-subtype reassortments is considered robust to detection bias.

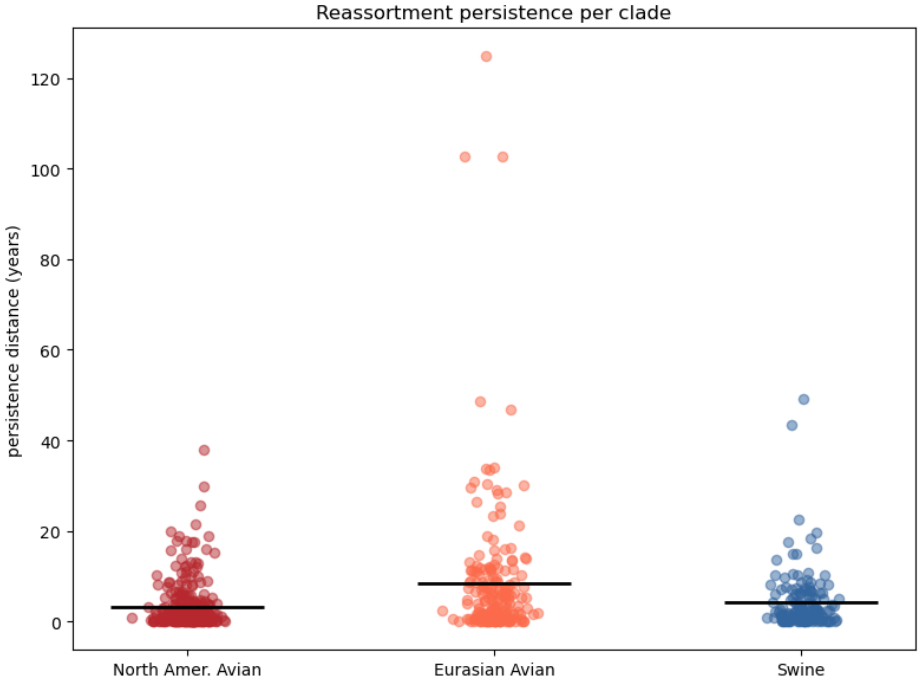


**Supplemental Figure 15: Average reassortment persistences are similar across host groups.** Persistence distance (years) was calculated for each detected reassortment event on an internal node across three host groups: North American avian (n = 306 lineages), Eurasian avian (n = 198), and swine (n = 150). Eurasian avian lineages showed the longest mean persistence (8.37 years, SD = 15.50, median = 3.49), followed by swine (mean = 4.22, SD = 6.60, median = 2.06), and North American avian (mean = 3.15, SD = 5.10, median = 1.22). Most lineages persist briefly with a subset persisting substantially longer in time. Horizontal bars indicate the mean. See methods for calculation details.

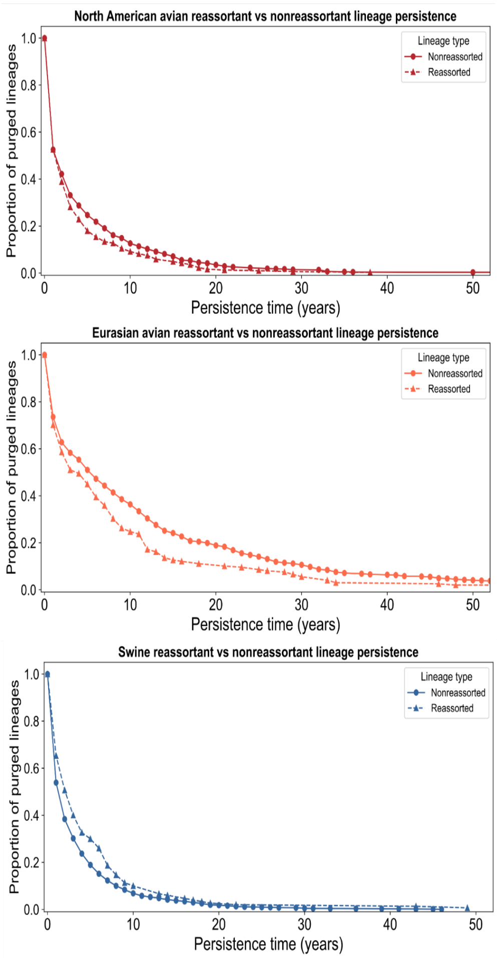


**Supplementary Figure 16. Reassortant and nonreassortant lineage persistence times by host group.** Proportion of reassortant lineages (dashed line, triangles) and nonreassortant (solid line, circles) lineages still in circulation at that time for North American avian, Eurasian avian, and swine host groups. Persistence time (x-axis) is defined as the maximum path length from a reassortant event to the nearest downstream reassortment event or terminal node, scaled by the host-specific molecular clock rate. Nonreassortant lineage persistences are defined similarly but starting with nodes lacking a high-support reassortment event.

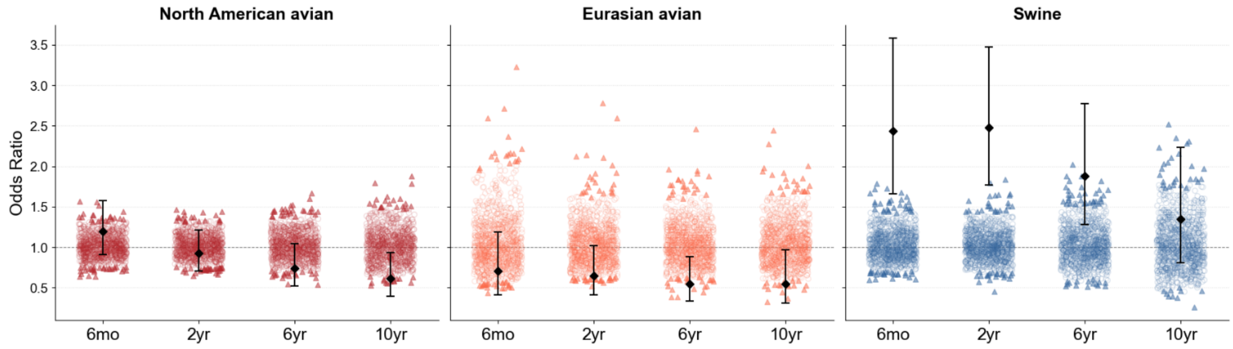


**Supplementary Figure 17. Fisher's exact test (FET) sensitivity analysis for reassortment-fitness associations.** For each host group, 1,000 null datasets were generated by randomly shuffling reassortment event positions across summary trees while preserving topology (See Methods for details). For each null tree, long-term survival effects of random reassortment were quantified using Fisher's exact test, where fitness was defined as a reassortment event sustaining descendants at x = 0.5, 2, 6, and 10 years into the future. Colored points represent the null distribution of odds ratios; triangles indicate null tests reaching statistical significance ($p < 0.05$) and circles indicate non-significant results. Black diamonds with error bars denote the observed odds ratio and 95% confidence interval.

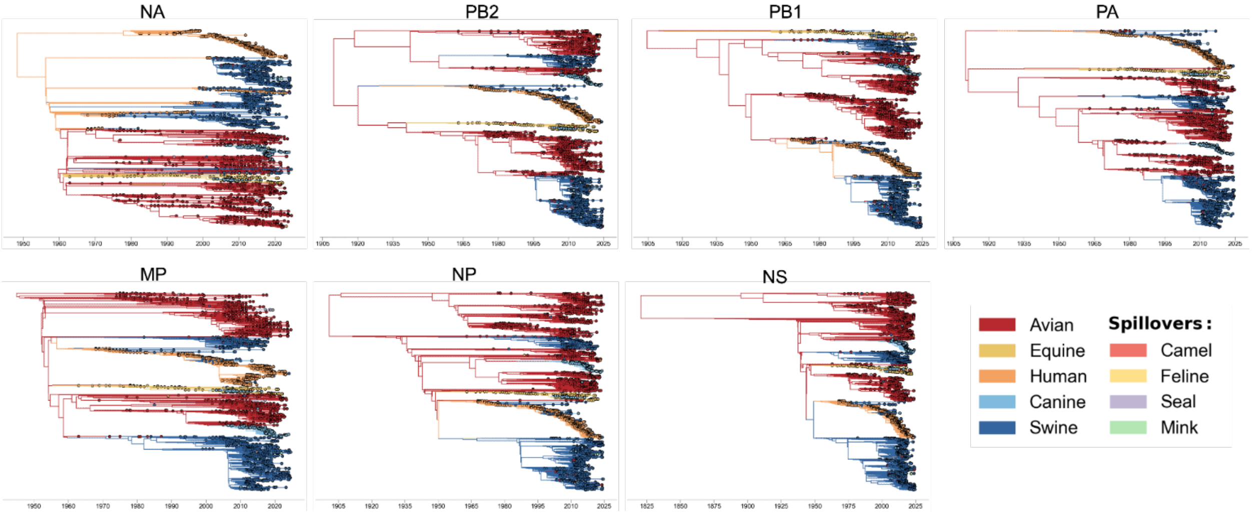


**Supplementary Figure 18: H3Nx viruses have jumped hosts multiple times.** The phylogenies for NA, PB1, PB1, PA, NP, NS, and MP (N=6,104) are colored by host type, where each leaf is a unique strain and each internal node is an inferred ancestor.

| Node name | Node type | Host switch | Segments |
|---|---|---|---|
| A/seal/Massachusetts/3911/1992\|1992-01-01 | leaf | avian --> seal | NS |
| TS_NODE_1376 | node | avian --> seal | PA |
| A/Phoca_vitulina/Alaska/PV2114/2021\|2021-08-24 | leaf | avian --> seal | NA |
| A/swine/Kazakhstan/106/1985\|1985-01-01 | leaf | avian --> swine | NA, NS |
| A/Changsha/1000/2022\|2022-05-11 | leaf | avian --> human | PB1 |
| A/Felis_catus/USA/047732/2018\|2018-03-27 | leaf | canine --> feline | MP, NP, NS, PA, PB1, PB2 |
| TS_NODE_6096 | node | avian --> human | NP |
| TS_NODE_3005 | node | human --> swine | MP, NP, NS, PA, PB2 |
| TS_NODE_3035 | node | human --> swine | NS |
| TS_NODE_3292 | node | human --> swine | NS |
| A/swine/Nagano/2000\|2000-01-01 | leaf | human --> swine | NA |
| A/swine/Hong_Kong/q066/99\|1999-01-01 | leaf | human --> swine | NA |
| TS_NODE_4651 | node | swine --> human | NA |
| TS_NODE_3656 | node | human --> swine | MP, NP, NS, PA, PB2 |
| TS_NODE_3921 | node | human --> swine | MP, NA, NP, NS, PA, PB1, PB2 |
| TS_NODE_3864 | node | swine --> avian | MP, NA, NP, PB1 |
| A/swine/Guatemala/CIP049-IP040078/2010\|2010-10-07 | leaf | human --> swine | NA |
| A/swine/Zambia/51/2018\|2018-09-01 | leaf | human --> swine | NA |
| TS_NODE_4109 | node | human --> swine | MP, NP, NS, PA, PB1, PB2 |
| TS_NODE_4192 | node | human --> swine | MP, NP, NS, PA, PB1, PB2 |
| A/chicken/Changzhou/c02/2013\|2014-04-17 | leaf | human --> avian | NA |
| TS_NODE_4318 | node | human --> swine | NA |
| A/swine/Guatemala/MM-160/2013\|2013-12-01 | leaf | human --> swine | NA, PA |
| TS_NODE_4467 | node | human --> swine | NA |
| TS_NODE_4588 | node | human --> swine | MP, NA, NP, NS, PA, PB2 |
| TS_NODE_4619 | node | human --> swine | NS |
| TS_NODE_4665 | node | human --> swine | MP, NP, PA, PB2 |
| TS_NODE_4724 | node | human --> swine | PA, PB1, PB2 |
| TS_NODE_5555 | node | human --> swine | MP, NA, NP, NS, PA, PB2 |
| A/mink/Wisconsin/31512-7/2012\|2012-08-14 | leaf | swine --> mink | NA |
| A/Iowa/08/2011\|2011-11-14 | leaf | swine --> human | NA |
| TS_NODE_5601 | node | human --> swine | MP, NA, NP, NS, PA, PB2 |
| TS_NODE_5661 | node | human --> swine | NA, NS |
| TS_NODE_6067 | node | human --> swine | MP, PA, PB2 |
| A/swine/Hong_Kong/126/1982\|1982-01-01 | leaf | human --> swine | NS |

**Supplementary Table 1. Reassortment events associated with host switching.** Each row represents a reassorted node or leaf tip in which the host annotation differs from that of its parent node, indicating a host switch coincident with reassortment. The segments column lists the segments inferred to have been reassorted in that event.
